# *SCN1A* haploinsufficiency disrupts neural progenitor networks and accelerates neurogenesis in human Dravet syndrome forebrain assembloids

**DOI:** 10.64898/2026.10.14.758626

**Authors:** Deepika Dogra, Harmony Fong, Sinan Zhang, Takuma Sumi, Nicole Rosin, Nicolas Collao, Marissa Gordon, Anneke Korver, Cezar Gavrilovici, Duncan Noble, Segundo Jose Guzman, Leili Rohani, Derrick Rancourt, Jeff Biernaskie, Julia Jacobs-Levan, Roger Thompson, Deborah M Kurrasch

## Abstract

Dravet syndrome is a severe childhood neurological disorder caused mostly due to mutations in the sodium channel gene *SCN1A*. Although the disorder is widely attributed to dysfunction of inhibitory neurons, growing evidence suggests that disease mechanisms may begin much earlier during brain development. To investigate the developmental consequences of *SCN1A* haploinsufficiency, we generated patient-derived dorsal-ventral forebrain assembloids that model key features of human corticogenesis. We found that SCN1A was expressed in ventricular zone neural progenitors and that its reduced expression was associated with impaired growth, disrupted ventricular zone organization, and progressive depletion of progenitor populations. We further demonstrated reduced progenitor proliferation and self-renewal, accelerated cell-cycle exit, and premature neuronal differentiation, leading to increased production of excitatory neurons and enhanced network excitability. Single cell transcriptomic analyses revealed reduced cycling radial glial cells, accelerated progenitor maturation, and dysregulation of gap junction and calcium signaling pathways. Following these findings, we examined progenitor calcium activity and found that *SCN1A* haploinsufficiency disrupted coordinated functional network organization, resulting in reduced recurrent connectivity and fragmentation of functional cellular assemblies. These abnormalities occurred despite preserved intrinsic electrophysiological properties of individual progenitors, indicating a defect in population-level coordination rather than single cell electrophysiology. Together, our findings identify disrupted progenitor communication as an early developmental consequence of *SCN1A* haploinsufficiency and suggest that abnormalities within neural progenitor populations contribute to altered corticogenesis, excitatory neurogenesis, and circuit hyperexcitability in Dravet syndrome. This work expands current disease models beyond interneuron dysfunction and highlights the developing ventricular zone as a potential therapeutic target.

**One-sentence summary:** *SCN1A* haploinsufficiency disrupts neural progenitor calcium network organization, impairing self-renewal and accelerating neurogenesis during human brain development.

## Introduction

Dravet syndrome (DS) is a severe developmental and epileptic encephalopathy characterized by pharmacoresistant seizures, intellectual disability, autism spectrum disorder-associated features, motor dysfunction, and an increased risk of sudden unexpected death in epilepsy (SUDEP) (1–5). More than 80% of DS cases are caused by heterozygous loss-of-function mutations in *SCN1A*, which encodes the voltage-gated sodium channel Nav1.1 (6, 7). Since the identification of *SCN1A* as the major disease-causing gene in DS, the disorder has largely been interpreted through the “interneuron hypothesis”, whereby reduced Nav1.1 activity impairs inhibitory GABAergic interneuron function, resulting in neural circuit hyperexcitability and seizures (8–11). Indeed, Nav1.1 is highly enriched within parvalbumin- and somatostatin-expressing interneurons, and impaired inhibitory neuron excitability has been consistently demonstrated in both animal and patient-derived models (8–12). However, the developmental origin of the symptoms observed in individuals with DS are only partially explained by interneuron dysfunction alone, suggesting that broader neurodevelopmental mechanisms may contribute to disease pathogenesis.

Increasing evidence suggests that *SCN1A* may function during brain development prior to the maturation of neuronal circuits. Transcriptomic studies and immunohistochemical analysis demonstrate that *SCN1A* expression is detectable in the human fetal brain and increases progressively throughout corticogenesis and postnatal development (13–15). Consistent with these observations, a recent preprint integrating human brain organoid models, mouse models, and postmortem tissue provided evidence that Nav1.1 may influence early corticogenesis (16), although the mechanisms through which *SCN1A* regulates neural progenitor behavior remain incompletely understood. Together, these findings challenge the traditional view of *SCN1A* as a regulator of mature neuronal excitability alone and suggest that this sodium channel may have critical developmental functions during early brain development.

The generation of the cerebral cortex involves coordinated regulation of neural progenitor populations residing within ventricular and subventricular germinal zones. Radial glial cells serve as the primary neural stem cells of the developing cortex, balancing self-renewal and differentiation to ensure appropriate cortical growth and neuronal production (17, 18). Perturbations of progenitor maintenance can lead to premature depletion of stem cell pools, altered neurogenesis, and abnormal cortical architecture, all of which have been implicated in neurodevelopmental disorders (19, 20). Importantly, neural progenitors do not function as isolated cells. Instead, radial glial cells communicate through gap junctions and coordinated calcium networks that regulate proliferation, differentiation, and tissue growth (21–25). Gap junction-dependent signaling has also been shown to influence the differentiation of human neural progenitor cells, highlighting the broader importance of coordinated progenitor communication during corticogenesis (25). Despite the central role of calcium network organization in regulating neural progenitor behavior, whether *SCN1A* contributes to these processes remains unknown.

Human induced pluripotent stem cell (iPSC)-derived brain organoids provide a powerful platform for investigating the developmental origins of neurological disorders (26–29). Forebrain assembloids generated through fusion of dorsal cortical organoids and ventral medial ganglionic eminence organoids recapitulate key aspects of human forebrain development, including ventricular zone formation, interneuron migration, and excitatory-inhibitory circuit assembly (30, 31). Although previous DS organoid studies have focused largely on neuronal excitability, interneuron dysfunction and metabolic perturbations (32–35), the mechanisms by which *SCN1A* haploinsufficiency influences neural progenitor development and corticogenesis remain poorly understood.

Here, we generated patient-derived forebrain assembloids to investigate the developmental consequences of *SCN1A* haploinsufficiency during human brain development. We show that SCN1A is expressed in ventricular zone neural progenitors and that its loss is associated with impaired progenitor maintenance, accelerated cell cycle exit, and premature neuronal differentiation. Single cell transcriptomic analyses revealed depletion of cycling radial glial cells accompanied by altered cell cycle dynamics, accelerated developmental maturation, and dysregulation of gap junction and calcium signaling pathways. Consistent with these findings, live calcium imaging identified disrupted progenitor functional network organization characterized by reduced recurrent coordination and fragmented coactive assemblies. Together, our findings identify a previously unrecognized developmental role for SCN1A in regulating progenitor network activity and suggest that early abnormalities in neural progenitor populations contribute to altered corticogenesis and neural circuit dysfunction in DS.

## Results

### Generation of patient-derived forebrain assembloids to model Dravet syndrome and expression analysis of SCN1A

To establish a human-relevant model of DS, peripheral blood mononuclear cells (PBMCs) obtained from a DS patient (hereafter referred to as DS) carrying a heterozygous de novo *SCN1A* missense variant, c.1876A>G (p.Ser626Gly), and her unaffected mother (hereafter referred to as control) were reprogrammed into iPSCs. These iPSCs were subsequently used to generate region-specific forebrain organoids that were fused to produce forebrain assembloids (Fig. 1A). Prior to differentiation, multiple iPSC clones were screened and validated for pluripotency. Both control and DS iPSC lines robustly expressed the canonical pluripotency markers OCT3/4, SOX2, SSEA-4, and TRA-1-60 (Fig. S1A). Furthermore, simple and cystic embryonic body (EB) aggregates were generated from these iPSCs and allowed to differentiate in culture to confirm their capacity to generate derivatives of all three germ layers (36), as demonstrated by the expression of α-fetoprotein (AFP; endoderm), βIII-tubulin (ectoderm), and α-smooth muscle actin (α-SMA; mesoderm) (Fig. S1B). These findings verified the pluripotent state and developmental competence of both control and DS iPSC lines.

**Figure 1.**
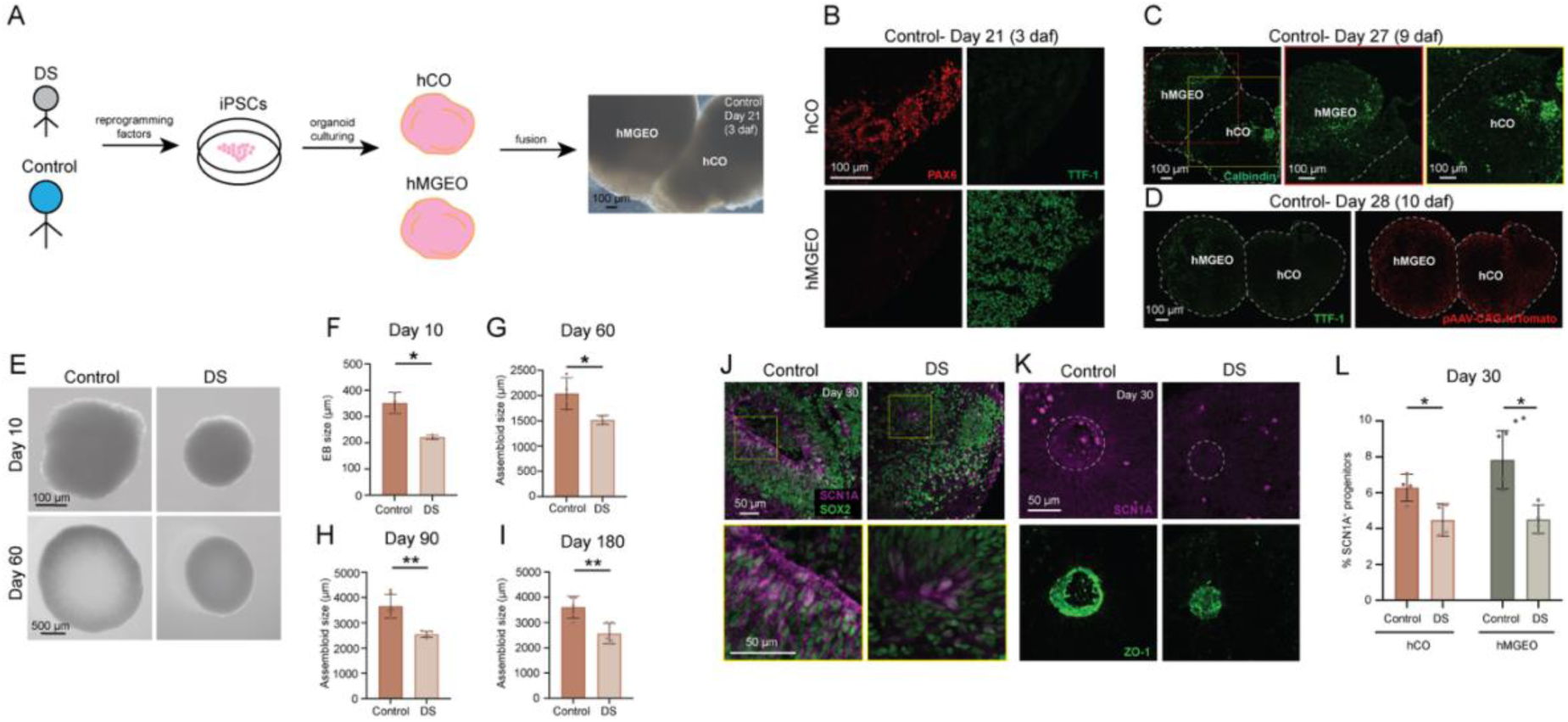
Generation and characterization of patient-derived forebrain assembloids reveals reduced growth and diminished progenitor SCN1A expression in DS. (A) Schematic overview of forebrain assembloid generation. PBMCs obtained from a DS patient carrying an *SCN1A* mutation and her healthy mother (control) were reprogrammed into iPSCs. iPSCs were differentiated into hCOs and hMGEOs, which were subsequently fused to generate forebrain assembloids. (B) Validation of dorsal-ventral regional specification in control assembloids at 3 daf (day 21). Immunostained sections demonstrate confinement of the dorsal forebrain progenitor marker PAX6 (red) to the hCO region and the ventral forebrain marker TTF-1 (NKX2.1; green) to the hMGEO region. Control, n=3. (C) Interneuron migration in control assembloids at 9 daf (day 27). Sections immunostained for Calbindin (green) identify migrating interneurons distributed within both hMGEO and hCO compartments. Control, n=3. (D) Validation of interneuron migration in control assembloids using AAV-CAG-tdTomato-labelled hMGEOs. At 10 daf (day 28), tdTomato^+^ cells (red) originating from the TTF-1^+^ (green) hMGEO region are observed migrating into the adjacent hCO region. Control, n=3. (E) Representative brightfield images of control and DS EBs (day 10) and forebrain assembloids (day 60), illustrating reduced growth of DS cultures. (F-I) Quantification of EB and assembloid size during development. DS cultures exhibit significantly reduced size relative to controls at day 10 (F), day 60 (G), day 90 (H) and day 180 (I). Control and DS, n=3 each genotype for day 10; n=4 each genotype for day 60; n=6 each genotype for day 90; n=5 each genotype for day 180. (J) Immunostaining of day 30 assembloid sections showing SCN1A expression (magenta) within SOX2+ (green) neural progenitors located in ventricular zones in control assembloids. DS assembloids exhibit reduced SCN1A expression within progenitor domains. Insets show higher magnification views of boxed regions. (K) Analysis of the localization of SCN1A at the apical ventricular surface on immunostained sections. In control assembloids, SCN1A (magenta) is enriched within ZO-1^+^ (green) ventricular zone structures, whereas this apically enriched signal is lost in DS assembloids. Control, n=3; DS, n=3. (L) Quantification of SCN1A^+^ progenitors within hCO and hMGEO regions at day 30. DS assembloids exhibit a significant reduction in the percentage of SCN1A^+^ progenitors (out of total SOX2^+^ progenitors) in both regions relative to controls. Control, n=4 for hCO and hMGEO; DS, n=4 for hCO and hMGEO. Data are Mean ± S.D., *P < 0.05, **P < 0.01-Welch’s t test. Each point on the graphs represents one assembloid. DS, Dravet syndrome; iPSCs, induced pluripotent stem cells; hCO, human cortical organoids; hMGEO, human median ganglionic eminence organoids; daf, days after fusion, EB, embryonic body.

Accurate modeling of human epilepsy requires both excitatory glutamatergic neurons generated in the dorsal forebrain and inhibitory GABAergic interneurons derived from the ventral forebrain that migrate into the cortex during development (37, 38). Therefore, we employed an established protocol (31) to generate dorsal forebrain organoids, referred to as human cortical organoids (hCOs), and ventral forebrain organoids, referred to as human medial ganglionic eminence organoids (hMGEOs). iPSCs were used to generate EBs by neural induction, and EBs were allowed to differentiate into cortical fate (hCOs) and MGE fate (hMGEOs). Following regional specification, hCOs and hMGEOs were fused to generate forebrain assembloids, enabling the migration of inhibitory interneurons from the ventral forebrain into the dorsal forebrain and the formation of integrated excitatory-inhibitory neural circuits (Fig. 1A, S2).

To confirm successful regional patterning following fusion, assembloids were examined three days after fusion (daf) using markers specific to dorsal and ventral forebrain identities. Immunostaining revealed that the dorsal progenitor marker PAX6 was restricted to the hCO region, whereas the ventral progenitor marker TTF-1 (NKX2.1) was confined to the hMGEO region (Fig. 1B). These results demonstrate that both domains retained their distinct regional identities after fusion. We next assessed interneuron migration, a hallmark feature of forebrain assembloid development. At 9 daf, immunostaining for the interneuron marker Calbindin revealed the presence of Calbindin^+^ cells within both the hMGEO and hCO regions, indicating successful migration of inhibitory interneurons from the ventral to the dorsal forebrain (Fig. 1C). To further validate this observation, hMGEOs were transduced with an AAV-CAG:tdTomato reporter prior to fusion with hCO. Analysis at 10 daf showed numerous tdTomato^+^ cells originating from the TTF-1^+^ hMGEO region and dispersing into the adjacent TTF-1-negative hCO region (Fig. 1D), further confirming robust interneuron migration across the fusion interface. Importantly, comparable patterns of regional marker expression and interneuron migration were observed in DS assembloids, with no clear differences in dorsal-ventral patterning or migratory behavior relative to controls (Fig. S3A, B). Together, these findings demonstrate the successful generation of patient-specific and control forebrain assembloids that recapitulate key developmental features of the human forebrain, including regional specification and interneuron migration. This model provides a robust platform for investigating how *SCN1A* mutations influence early neurodevelopmental processes and contribute to DS pathogenesis.

Although *SCN1A*-related DS is not typically associated with a global change in the brain size at birth or early childhood, patients and animal models can develop region-specific brain atrophy and structural abnormalities over time (7, 39–41). Intriguingly, we observed that the DS assembloids were significantly smaller than controls throughout the in vitro development, quantified as early as day 10 to day 180 (−37.1% day 10; −25.6% day 60; −30.2% day 90; −28.3% day 180; Fig. 1E-I), signifying possible developmental defects due to *SCN1A* haploinsufficiency. To validate our findings, we generated assembloids from an independent DS patient-derived iPSC line (DD1C; referred to as DS_2 here) (42) and compared them with assembloids generated from an unrelated healthy control iPSC line (ACS-1023; referred to as control_2). Consistent with our initial observations, DS_2 assembloids were significantly smaller than control_2 EBs at day 10 and day 60 (−40.3% day 10; −24% day 60; Fig. S4), providing independent evidence that *SCN1A* haploinsufficiency negatively affects brain development.

SCN1A is widely expressed across different brain regions, with particularly high expression in the cerebral cortex, hippocampus, cerebellum, and other forebrain regions. In both humans and rodents, SCN1A expression is shown to be enriched in GABAergic interneurons, especially parvalbumin^+^ and somatostatin^+^ interneuron populations (3, 8, 13). Recent Scn1a-GFP reporter mouse work further refined this expression pattern by showing that while SCN1A is predominantly expressed in inhibitory neurons throughout the neocortex, it is also present in specific excitatory projection neuron populations, including most layer V pyramidal tract neurons and subsets of layer II/III corticocortical neurons (43). Published work and publicly available transcriptomic datasets (Fig. S5) have shown that SCN1A expression is detectable during human fetal brain development and increases progressively throughout postnatal neuronal maturation and circuit formation (14, 15, 44). Moreover, a recent preprint from Lignani et al. suggests that SCN1A is expressed during early corticogenesis in a human brain organoid model (16).

In line with published work, immunostaining of mature control assembloids at day 90 demonstrated SCN1A expression in both excitatory and inhibitory neurons (Fig. S6). We next sought to determine whether SCN1A is also expressed during early stages of development and whether reduced assembloid growth could be linked to impaired SCN1A function in neural progenitors. Immunostaining of day 30 control assembloids revealed clear SCN1A expression within SOX2^+^ neural progenitor cells located in ventricular zone-like structures (Fig. 1J). Notably, SCN1A expression appeared enriched at the apical surface of these ventricular zones, as demonstrated by co-localization with the apical junction marker ZO-1 (Fig. 1K). In contrast, DS assembloids exhibited a significant reduction in SCN1A expression within SOX2^+^ progenitors in both the hCO and hMGEO regions (−28.7% hCO; −42.3% hMGEO; Fig. 1J, L). Moreover, the apically enriched SCN1A signal observed in control ventricular zones was largely absent in DS assembloids (Fig. 1K), indicating a loss of SCN1A expression at the ZO-1^+^ ventricular surface. These data suggest that SCN1A is expressed in the neural progenitors within the ventricular zones of the developing cortex and that *SCN1A* haploinsufficiency may contribute to the developmental deficits underlying the reduced size of DS assembloids.

### Neural progenitor self-renewal is negatively affected in DS assembloids

Given the expression of SCN1A within ventricular zone progenitors and the reduced growth of DS assembloids, we next investigated whether *SCN1A* haploinsufficiency disrupts progenitor maintenance. Hoechst staining at day 30 revealed that defects in SCN1A function not only affected the assembloid size but also caused compromised ventricular zone architecture as shown by comparing the whole assembloid structure between DS and control, as well as DS_2 and control_2 assembloids (Fig. S7). Further, by performing immunostaining for SOX2, we quantified the neural progenitor population at day 30, day 60 and day 90. Our DS assembloids showed a significant reduction in the percentage of SOX2^+^ cells in both the hCO and hMGEO regions at all the stages of development relative to controls (−29% hCO and −30% hMGEO day 30; −46% hCO and −40% hMGEO day 60; −37.6% hCO and −28.1% hMGEO day 90; Fig. 2A-D). This finding was independently validated using the other DS iPSC line (DS_2), which similarly exhibited a significant reduction in the percentage of SOX2^+^ cell population in both the hCO and hMGEO regions compared to control_2 assembloids at day 60 (−37.7% hCO; −42.9% hMGEO; Fig. S8). To determine how the depletion of neural progenitors affected ventricular zone architecture, we quantified both the number and thickness of ventricular zones in SOX2-immunostained assembloids. The number of SOX2^+^ ventricular zones was significantly reduced across development in the DS assembloids compared to the control assembloids (−43% day 30; −74.6% day 60; −82.5% day 90; Fig. S9A-D). This was accompanied by progressive thinning of ventricular zone structures at later stages in the DS assembloids relative to control assembloids (−33.2% day 60; − 45.1% day 90; Fig. S9A, E-G). Collectively, these findings indicate that *SCN1A* haploinsufficiency compromises the maintenance of neural progenitor populations, resulting in the disruption of ventricular zone architecture during brain development. Such defects may contribute to the reduced growth phenotype observed in DS assembloids. Our findings are further supported by previous observations made in a mouse DS model including neurogenic niche alterations with fewer neural stem cells and impaired cell division, and a human cortical organoid DS model showing alterations in neural progenitor cell cycle progression (16, 45).

**Figure 2.**
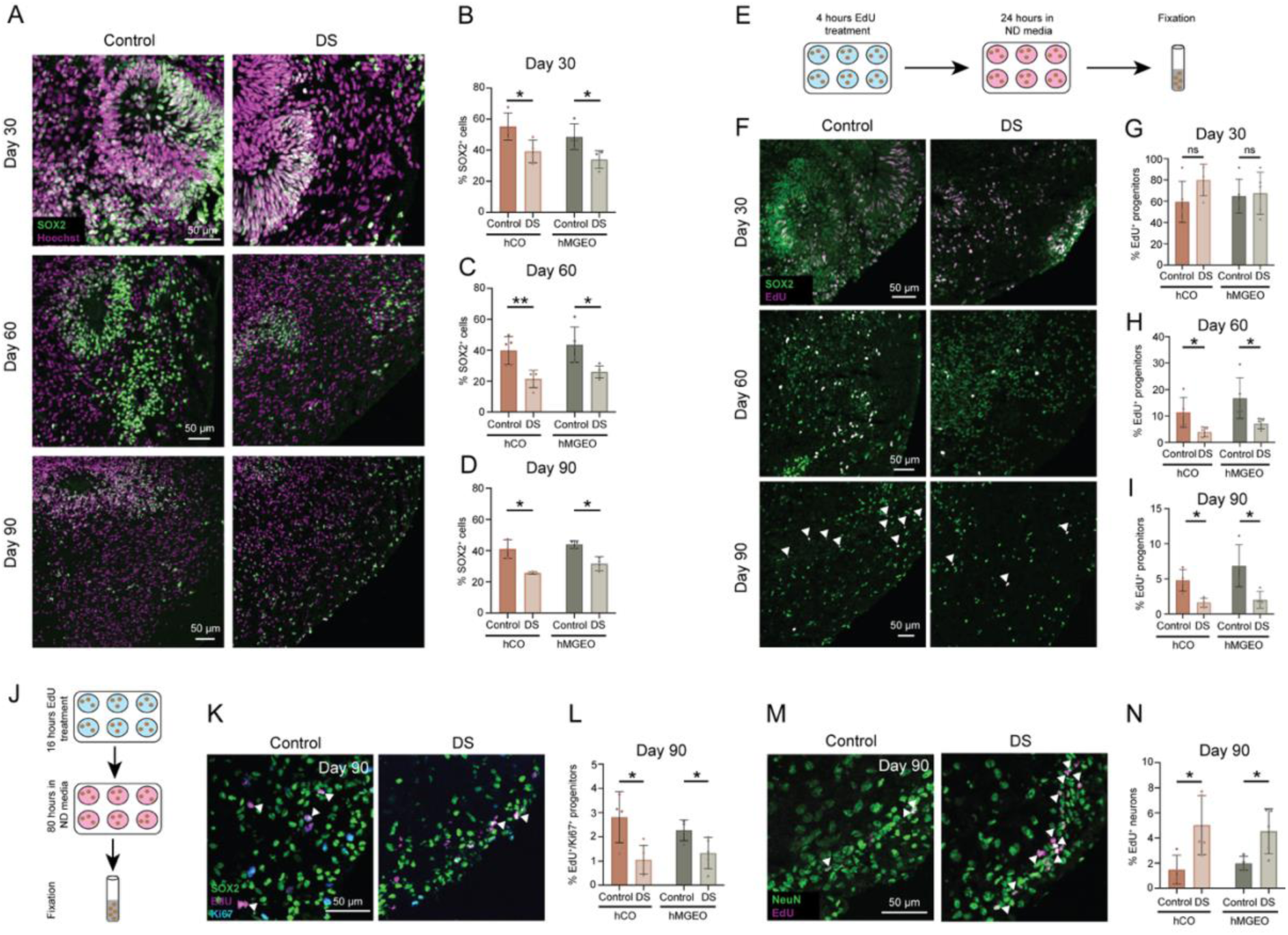
*SCN1A* haploinsufficiency impairs neural progenitor maintenance, proliferation and self-renewal in forebrain assembloids. (A) Representative immunofluorescence images of day 30, day 60 and day 90 control and DS assembloid sections stained for the neural progenitor marker SOX2 (green) and Hoechst (magenta). DS assembloids exhibit a progressive reduction in SOX2^+^ progenitor populations relative to controls. (B-D) Quantification of SOX2^+^ neural progenitors in the hCO and hMGEO regions at day 30 (B), day 60 (C) and day 90 (D). DS assembloids show significantly reduced percentage of SOX2^+^ cells (out of total Hoechst^+^ cells) across development. Control and DS, n=4 each genotype for hCO and hMGEO at day 30; n=5 each genotype for hCO and hMGEO at day 60; n=3 each genotype for hCO and hMGEO at day 90. (E) Experimental design for short-term EdU pulse-chase analysis. Assembloids were exposed to EdU for 4 hours, followed by a 24-hour chase period in ND media prior to fixation and analysis. (F) Representative images of control and DS assembloids at day 30, day 60 and day 90 following EdU pulse labelling. Sections were immunostained for SOX2 (green) and EdU (magenta) to identify proliferating neural progenitors. Arrowheads indicate SOX2^+^/EdU^+^ cells. (G-I) Quantification of EdU^+^ progenitors in the hCO and hMGEO regions at day 30 (G), day 60 (H) and day 90 (I). No differences were observed at day 30, whereas DS assembloids exhibited a significant reduction in the percentage of EdU^+^ progenitors (out of total SOX2^+^ progenitors) at later developmental stages. Control and DS, n=5 each genotype for hCO and hMGEO at day 30; n=5 each genotype for hCO and hMGEO at day 60; n=4 each genotype for hCO and hMGEO at day 90. (J) Experimental design for long-term EdU pulse-chase analysis used to assess neural progenitor self-renewal and cell cycle exit. Day 90 assembloids were treated with EdU for 16 hours, followed by an 80-hour chase period in ND media prior to fixation. (K) Representative images of day 90 control and DS assembloids immunostained for SOX2 (green), Ki67 (cyan) and EdU (magenta). Arrowheads indicate SOX2^+^/Ki67^+^/EdU^+^ cells. (L) Quantification of EdU^+^/Ki67^+^ progenitors in the hCO and hMGEO regions at day 90. DS assembloids exhibit a reduced proliferative progenitor fraction (percentage of EdU^+^/Ki67^+^ progenitors out of total SOX2^+^ progenitors) indicative of impaired self-renewal. Control, n=4 for hCO and hMGEO; DS, n=5 for hCO and hMGEO. (M) Representative images of day 90 control and DS assembloids immunostained for NeuN (green) and EdU (magenta). Arrowheads indicate NeuN^+^/EdU^+^ cells. (N) Quantification of EdU^+^ neurons in the hCO and hMGEO regions at day 90. DS assembloids exhibit an increased quiescent fraction (percentage of EdU^+^ neurons out of total NeuN^+^ neurons), consistent with accelerated cell cycle exit and premature neuronal differentiation. Control, n=4 for hCO and hMGEO; DS, n=5 for hCO and hMGEO. Data are Mean ± S.D., ns: no significant changes observed, *P < 0.05, **P < 0.01-Welch’s t test. Each point on the graphs represents one assembloid.

To determine whether progenitor depletion resulted from increased cell death, we performed co-immunostaining for SOX2 and cleaved caspase-3 (CC3; apoptosis marker) at day 30 and day 90, quantifying progenitor apoptosis. No significant changes were observed between control and DS assembloids (Fig. S10A-D), indicating that enhanced apoptosis is unlikely to account for progenitor loss. We next questioned whether defective cell proliferation was a potential cause of reduction in progenitor population. We treated assembloids with EdU by adding it to the neural differentiation (ND) media for 4 hours followed by washing out EdU and leaving them in ND media alone for 24 hours (Fig. 2E). Later, the assembloids were fixed, sectioned, immunostained for SOX2, and labelled with EdU to quantify dividing progenitors. Although no differences were detected at day 30, both the hCO and hMGEO regions of the DS assembloids showed a significant reduction in the percentage of EdU^+^ progenitors relative to controls at day 60 and day 90 (−65.2% hCO and −57.7% hMGEO day 60; −66.2% hCO and −70.5% hMGEO day 90; Fig. 2F-I), demonstrating that the decline in the progenitor population is possibly a consequence of a robust deficit in their proliferative capacity.

Next, to determine the effect of compromised progenitor proliferation on their self-renewal and subsequent neurogenesis, we measured their cell cycle kinetics and estimated the proportion of proliferative (P) and quiescent (Q) fractions using EdU labelling. We treated the assembloids at day 90 with EdU (added to the ND media) for 16 hours, washed out EdU, kept the assembloids in ND media alone for 80 hours to allow EdU to label the dividing cells, followed by their fixation and sectioning (Fig. 2J). We co-immunostained for SOX2 and Ki67, followed by EdU labelling, and observed a significant reduction in the percentage of Ki67^+^/EdU^+^ progenitors in both the hCO and hMGEO regions of the DS assembloids compared to controls (−62.3% hCO; −41.4% hMGEO; Fig. 2K, L), indicative of a decline in the progenitor P fraction index. Simultaneously, we immunostained for the neuron marker NeuN and labelled with EdU. There was a significant increase in the percentage of EdU^+^ neurons in both the hCO and hMGEO regions of the DS assembloids compared to control ones (241.5% hCO; 128.1% hMGEO; Fig. 2M, N), indicating an accelerated Q fraction index of the progenitors. Collectively, these data demonstrate that *SCN1A* haploinsufficiency impairs neural progenitor self-renewal and promotes premature cell cycle exit to give rise to neurons. This progenitor imbalance likely contributes to ventricular zone depletion and the reduced size phenotype observed in DS assembloids.

### *SCN1A* haploinsufficiency drives premature neurogenesis towards excitatory neurons and causes hyperexcitability

Given that neural progenitors in DS assembloids exited the cell cycle more rapidly and generated neurons prematurely, we next assessed whether overall neuronal populations were altered. Thus, we immunostained assembloid sections for the neuron marker NeuN at day 60 and day 90, and quantified the neuron population. There was a significant increase in the percentage of NeuN^+^ cells in both the hCO and hMGEO regions of the DS assembloids compared to controls (135.1% hCO and 93.3% hMGEO day 60; 91.7% hCO and 137.1% hMGEO day 90; Fig. 3A-C),, consistent with accelerated neurogenesis. These data were further validated using the second DS iPSC line (DS_2), which similarly exhibited a significant increase in the percentage of NeuN^+^ cell population in both the hCO and hMGEO regions compared to control_2 assembloids at day 60 (83.47% hCO and 77.64% hMGEO; Fig. S11).

**Figure 3.**
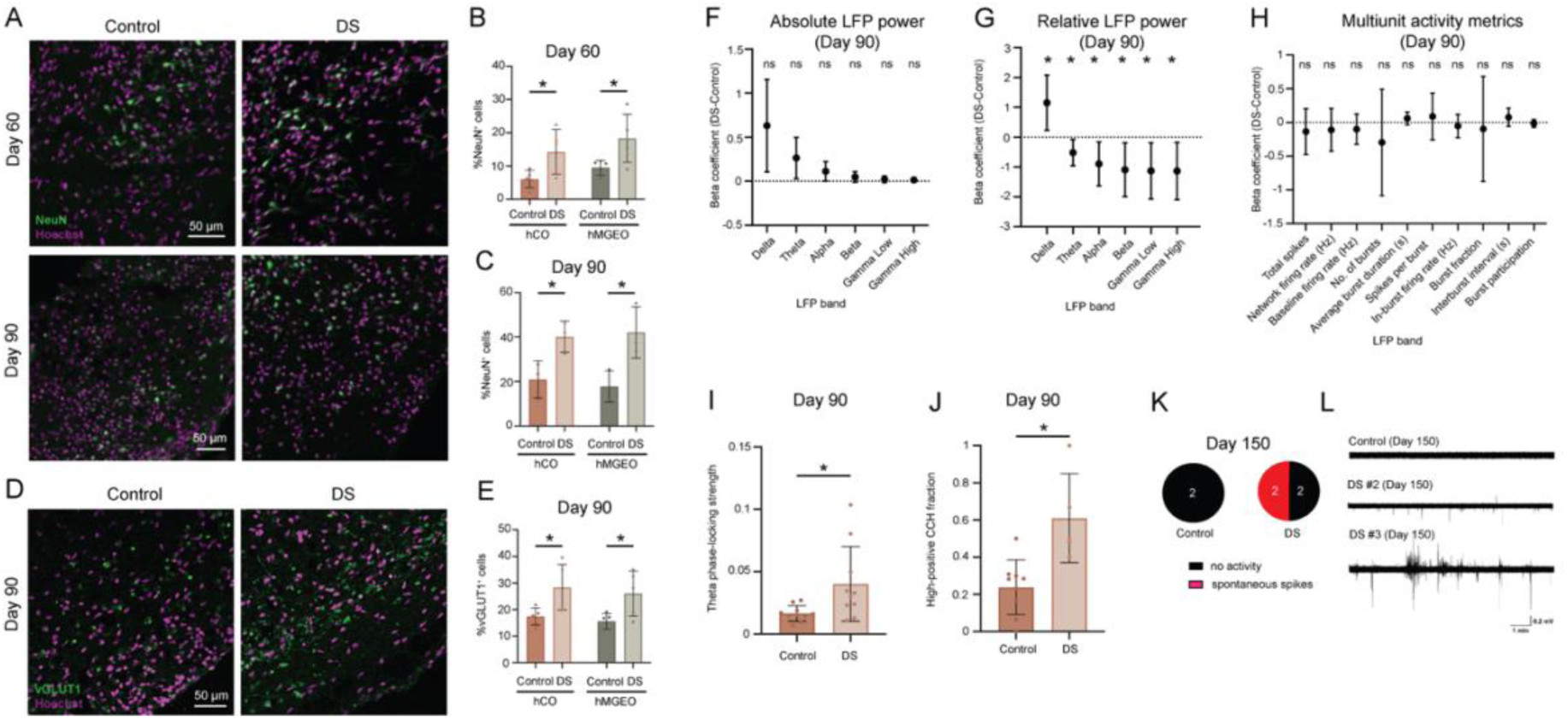
*SCN1A* haploinsufficiency drives premature excitatory neurogenesis and promotes hyperexcitable neural network activity. (A) Representative images of control and DS assembloids at day 60 and day 90 immunostained for NeuN (green) and Hoechst (magenta). DS assembloids exhibit an increased neuronal population relative to controls. (B-C) Quantification of NeuN^+^ neurons in the hCO and hMGEO regions at day 60 (B) and day 90 (C). DS assembloids show a significant increase in the percentage of NeuN^+^ cells (out of total Hoechst^+^ cells) in both forebrain compartments compared to control ones. Control and DS, n=5 each genotype for hCO and hMGEO at day 60; n=3 each genotype for hCO and hMGEO at day 90. (D) Representative images of day 90 control and DS assembloids immunostained for vGLUT1 (green) and Hoechst (magenta). DS assembloids display an increased abundance of vGLUT1^+^ cells. (E) Quantification of vGLUT1^+^ excitatory neurons within the hCO and hMGEO regions at day 90. The percentage of vGLUT1^+^ cells (out of total Hoechst^+^ cells) is significantly increased in DS assembloids relative to controls, indicating enhanced excitatory neurogenesis associated with *SCN1A* haploinsufficiency. Control, n=5 for hCO and hMGEO; DS, n=5 for hCO and hMGEO. Data are Mean ± S.D., *P < 0.05-Welch’s t test for panels A-E, J. Each point on the graphs represents one assembloid. (F, G) Electrophysiological analyses at day 90. Estimated group effects for absolute (F) and relative (G) LFP power across frequency bands. Absolute power did not differ significantly between groups (ns: no significant changes, Q > 0.05 after FDR correction), whereas DS assembloids showed significantly increased relative delta power and reduced relative power in all other frequency bands (*Q < 0.05 after FDR correction). Frequency bands were defined as: delta, 1–4 Hz; theta, 4–8 Hz; alpha, 8–12 Hz; beta, 12–30 Hz; low-gamma, 30–55 Hz; and high-gamma, 65–100 Hz. (H) Estimated group effects for MUA measures. No MUA metric differed significantly between groups after FDR correction (ns: no significant changes, Q > 0.05). Points represent estimated DS minus control effects (beta coefficients), and error bars represent 95% confidence intervals; the dotted line indicates no group difference. For panels F-H, group differences were assessed using linear mixed-effects models with group as a fixed effect and assembloid identity as a random intercept. When model fitting failed or yielded singular fits, Welch’s t-tests were performed on assembloid-level means. (I) Theta-band MUA-LFP phase-locking strength. DS assembloids showed greater theta phase-locking strength than controls. Data are Mean ± S.D., *P < 0.05-Welch’s t test. Control, n=10; DS, n=10 for panels F-I. (J) High-positive CCH fraction analysis showing a greater prevalence of strongly positive correlations in DS assembloids. Control, n = 9; DS, n = 5. Data are Mean ± S.E.M., *P < 0.05-Welch’s t test. (K) A pie chart of the number of control and DS assembloids showing different patterns of extracellular field potentials at day 150. 50% of the DS assembloids show spontaneous spikes unlike control that show no abnormal activity. Control, n=2; DS, n=4. (L) Representative extracellular field recordings performed at day 150. Traces are 10-min recordings from the most active site sampled in each assembloid. Spontaneous spikes are visible in DS assembloid #2 and #3; the remaining recordings show no detectable events. LFP, local field potential; MUA, multiunit activity; CCH, cross-correlation histograms.

To determine whether specific neuronal subtypes were differentially affected by *SCN1A* haploinsufficiency, we immunostained assembloid sections for the excitatory neuron marker vGLUT1 and inhibitory neuron marker GABA at day 90, and quantified the two neuronal populations. The percentage of vGLUT1^+^ cells in both the hCO and hMGEO regions of the DS assembloids were significantly higher relative to the control assembloids (Fig. 3D, E); however the percentage of GABA^+^ cells were unchanged (Fig. S12A, B). Together, these findings indicate that *SCN1A* haploinsufficiency promotes premature neuronal differentiation that is biased towards an excitatory neuronal fate, while inhibitory neuron generation appears largely unaffected. This observation is particularly notable given previous studies demonstrating impairments in inhibitory neuron development and function, including disrupted interneuron maturation and altered action potential firing, in DS patient-derived models and individuals with DS (12, 34, 46). Our results therefore suggest that early neurodevelopmental consequences of *SCN1A* haploinsufficiency may include enhanced excitatory neurogenesis, potentially contributing to altered excitation-inhibition balance through mechanisms distinct from those previously described.

Next, we sought to investigate whether an enhanced population of excitatory neurons is causing disruptions in the neural activity of the DS assembloids. Using NeuroProbe, we recorded extracellular field potential of assembloids at day 90. LFP (local field potential) and MUA (multiunit activity) analyses included 20 assembloids (10 control and 10 DS) and 60 recordings in total. Absolute LFP power was unchanged in the primary analysis (Fig. 3F). In contrast, DS assembloids showed significantly increased relative delta power (67.8%; beta = 1.15, 95% CI 0.23 to 2.08, q = 0.022) and reduced relative power in other frequency bands relative to controls (Fig. 3G). Increase in the slow-frequency delta power is indicative of increased network synchronization during epilepsy (47–51) that aligns with our observation of an increase in excitatory neuron composition in the DS assembloids. Similarly, interburst analyses showed a similar redistribution, with increased absolute and relative delta power and reduced relative alpha, beta, low-gamma, and high-gamma power (Fig. S13A, B). Network-level MUA and burst metrics did not differ between groups after FDR correction (Fig. 3H).

*SCN1A* haploinsufficiency led to a 135.3% increase in theta LFP-MUA phase-locking strength in DS assembloids compared to controls (Fig. 3I; beta = 0.78, q = 0.023; Fig. 3I), further suggestive of enhanced network synchronization (52, 53). In contrast, high-gamma coherence was similar between groups, with the DS assembloids showing slightly lower pairwise coherence (Fig. S14B). Coherence decreased modestly with interelectrode distance, but neither the group effect nor the group-by-distance interaction was significant (Fig. S14C).

Spike-sorting analyses incorporated the same 20-assembloid, 60-recording cohort and yielded 151 curated units (an example shown in Fig S15A). At least one curated unit was detected in 43 recordings from 18 assembloids. Two DS assembloids yielded no curated units and therefore did not contribute to unit-level summary metrics. DS assembloids showed a lower within-unit burst fraction than controls (beta = −2.32 on the logit scale, 95% CI −3.87 to −0.77, q = 0.0299; Fig. S15B). Curated-unit yield, active-channel fraction, sorted-population firing rate, spike count, mean firing rate, ISI variability, waveform amplitude, trough-to-peak duration, and template spatial spread did not differ significantly after FDR correction (Fig. S15B-D). The increase in phase locking was further supported by pairwise spike-train cross-correlation analysis, which revealed a greater prevalence of strongly positive correlations in DS assembloids compared to controls (154% increase; Fig. 3J), without significant group differences in the mean positive peak or negative trough magnitude (Fig. S15E-G).

To further test the electrical activity of the assembloids at a later stage of development, we performed extracellular field recordings at day 150 on a small cohort of samples. We observed no spontaneous activity in either of the two control assembloids; however, two out of four DS assembloids exhibited spontaneous spiking (small amplitude and low frequency spikes; Fig. 3K, L). Specifically, DS assembloid #2 showed sparse spikes (amplitude = 0.29 ± 0.06 mV; frequency = 0.016 Hz) and DS assembloid #3 showed more frequent events (amplitude = 0.27 ± 0.07 mV; frequency = 0.22 Hz).

Together, these electrophysiological findings support increased network excitability in DS assembloids associated with *SCN1A* haploinsufficiency-induced excitatory neurogenesis. DS assembloids exhibited increased slow-frequency synchronization and enhanced theta phase-locking at day 90, while spontaneous spiking activity was detected in a subset of DS assembloids, but not in the control assembloids examined, at day 150. Collectively, these data suggest that excess excitatory neuron production contributes to persistent alterations in neural network activity and may promote the emergence of seizure-like phenotypes in DS neural circuits.

### scRNA-seq reveals that *SCN1A* haploinsufficiency accelerates neural progenitor maturation towards neuronal fate

To investigate the cellular mechanisms underlying the premature neurogenesis observed in DS assembloids, we performed single cell RNA sequencing (scRNA-seq) on pooled assembloids obtained from controls (control: mother and control_3: father) and DS assembloids (DS: daughter and DS_2: unrelated patient) at day 90. We identified multiple neural progenitor and differentiated cell populations, including cycling radial glial cells (RGCs), RGCs, intermediate progenitor cells (IPCs), excitatory neurons, inhibitory neurons, astroglia, and choroid plexus cells (Fig. 4A, S16, S17). Comparison of cell type composition across samples revealed shifts in progenitor populations, especially a decline in the proportion of cycling RGCs in DS assembloids relative to controls, suggesting altered developmental progression within early neural lineages (Fig. 4B).

**Figure 4.**
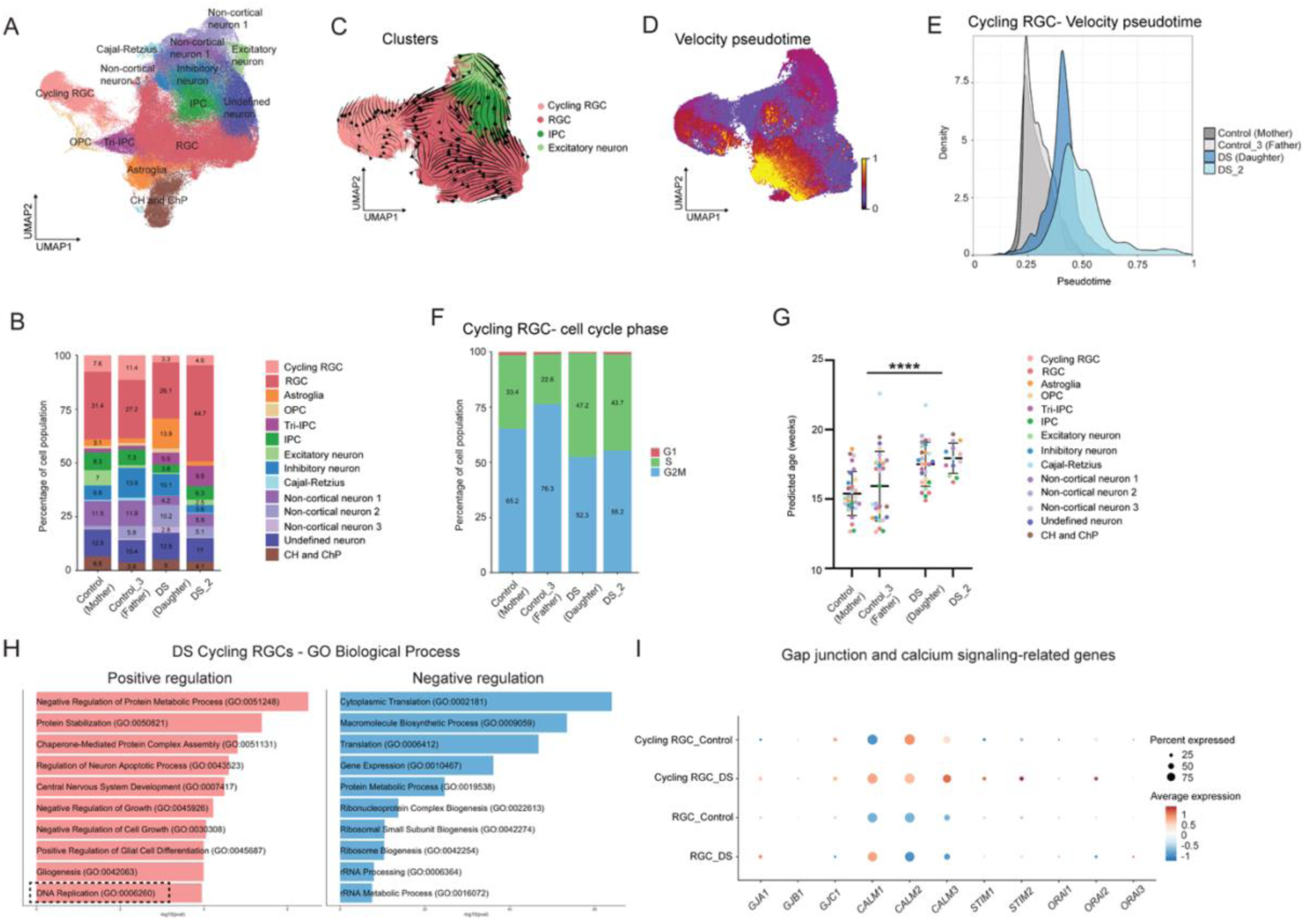
*SCN1A* haploinsufficiency reduces cycling radial glial cell abundance and accelerates progenitor maturation through altered cell cycle and signaling programs. (A) Integrated UMAP plot of neural cell types in control and DS assembloids. Control = 2 replicates, control_3 = 2 replicates, DS = 2 replicates, DS_2 = 1 replicate. (B) Stacked bar plot showing cell type composition across samples with a reduction in the proportion of cycling RGCs in the DS assembloids compared to controls. (C) UMAP plot of subsetted excitatory lineage cells (cycling RGCs, RGCs, IPCs, and excitatory neurons) with overlaid RNA velocity streamlines depicting developmental trajectories. (D) UMAP plot of subsetted excitatory lineage cells coloured by RNA velocity-inferred pseudotime values. (E) Density plot displaying RNA velocity-inferred pseudotime values of cycling RGCs. DS assembloids seem to occupy more advanced pseudotime states than controls. (F) Stacked bar plot showing cell cycle phase composition of cycling RGCs. DS cycling RGCs show a shift in cell cycle state relative to controls. (G) Predicted developmental age of cell types across control and DS samples using a cell type-agnostic model. DS assembloids display older predicated ages compared to controls (compared as control vs. DS group). Control = 2 replicates, control_3 = 2 replicates, DS = 2 replicates, DS_2 = 1 replicate. Control group, n=28 for control and n=28 for control_3; DS group, n=28 for DS and n=13 for DS_2. Data are Mean ± S.D., ****P < 0.0001-Welch’s t test. Each point on the graph represents one cell type. (H) GO enrichment analysis depicting the top 10 positively (red) and negatively (blue) regulated biological processes in DS cycling RGCs compared to control. DS assembloids show an upregulation of DNA replication process. (I) Dot plot showing relative expression of gap junction and calcium signaling-related genes in progenitor populations across combined control and DS groups. DS assembloids show dysregulated expression of some gap junction and calcium signaling-related genes. CH, cortical hem; ChP, choroid plexus; IPC, intermediate progenitor cell; OPC, oligodendrocyte precursor cell; RGC, radial glial cell; GO, Gene ontology.

Given our previous findings of accelerated cell cycle exit and premature neuronal differentiation, we next examined developmental trajectories within the progenitor compartment. RNA velocity analysis identified developmental trajectories originating from cycling RGCs and progressing towards more differentiated states, as expected (Fig. 4C). Strikingly, computation of pseudotime values based on splicing kinetics revealed that cycling RGCs in the DS assembloids occupied more advanced pseudotime states than control assembloids, consistent with accelerated developmental progression towards neuronal lineages (Fig. 4D, E).

To determine whether altered cell cycle dynamics accompanied this accelerated maturation, we examined cell cycle phase distributions within cycling RGCs. DS cycling RGCs exhibited a marked shift in cell cycle state relative to controls, characterized by an increased proportion of cells in S phase and a corresponding reduction in G2/M phase cells (Fig. 4F). This accumulation of progenitors in S phase suggests altered cell cycle progression in DS assembloids. Consistent with this observation, developmental age prediction using a published cell type-agnostic model (54) revealed that DS assembloids displayed significantly older predicted ages than control assembloids (Fig. 4G). Together, these findings indicate that *SCN1A* haploinsufficiency primarily impacts early progenitor populations, accelerating their progression towards more mature developmental states despite altered cell cycle dynamics.

To identify molecular pathways affected due to *SCN1A* haploinsufficiency, we performed gene ontology enrichment analysis of differentially expressed genes (DEGs) in DS cycling RGCs. Upregulated genes were enriched for biological processes related to protein metabolism, protein localization, neuronal development, cell growth, glial differentiation, and notably, DNA replication (Fig. 4H). The enrichment of DNA replication pathways is consistent with the increased accumulation of DS progenitors in S phase, further supporting the presence of dysregulated cell cycle progression within this population. In contrast, downregulated genes were enriched for pathways involved in cytoplasmic translation, ribosome biogenesis and RNA metabolism (Fig. 4H).

Because coordinated communication between neural progenitors is critical for maintaining appropriate developmental timing, we next examined gap junction genes. DS cycling RGCs exhibited altered expression of multiple connexin genes, including *GJA1*, *GJB1*, and *GJC1* (Fig. 4I). Gap junction-mediated coupling plays an important role in coordinating progenitor behavior during corticogenesis, and connexin proteins such as Cx43 (encoded by *GJA1*) have been implicated in regulating neural progenitor differentiation and fate specification (23, 25, 55). The dysregulation of multiple connexin genes in DS progenitors therefore suggests impaired intercellular communication within the progenitor niche. In parallel, several calcium-responsive genes, including *CALM1*, *CALM3*, *STIM1*, *STIM2*, *ORAI1*, and *ORAI2*, were upregulated in DS assembloids (Fig. 4I), indicating altered calcium signaling. Calcium transients and connexin-dependent calcium waves regulate radial glial proliferation and neurogenesis during cortical development (22, 56, 57), while Cx43 regulates differentiation of human neural progenitor cells (25). Accordingly, the concurrent dysregulation of connexin and calcium-responsive genes suggests that *SCN1A* haploinsufficiency may perturb signaling pathways that coordinate progenitor maturation and neurogenic timing.

Together, these results demonstrate that *SCN1A* haploinsufficiency disrupts neural progenitor cell cycle dynamics while accelerating progression towards more mature developmental states. DS progenitors exhibited advanced pseudotime positioning, higher predicted developmental age, accumulation within S phase, and transcriptional signatures associated with enhanced DNA replication, altered gap junction coupling, and dysregulated calcium-responsive signaling pathways. Overall, these findings identify aberrant progenitor maturation as a potential mechanism underlying the premature neurogenesis observed in DS assembloids and suggest that disrupted progenitor communication and calcium-dependent signaling contribute to altered neurodevelopmental trajectories in DS.

### *SCN1A* haploinsufficiency alters progenitor calcium network dynamics and promotes fragmented coactive assemblies

Because scRNA-seq analysis identified dysregulation of multiple connexin genes and calcium-responsive signaling pathways in DS cycling radial glia, we next tested whether these transcriptional alterations were accompanied by functional changes in progenitor calcium network activity. We performed live calcium imaging in the Fluo-4 AM labelled assembloids at day 30 (Movies S1, S2), a developmental stage enriched in ventricular zone-like progenitors and prior to the emergence of mature neuronal activity. We have shown that there are <10% Neun^+^ neurons and >50% SOX2^+^ neural progenitors at day 30 in the control assembloids (Fig. S18A, B), indicating that the majority of Fluo-4 AM-labeled cells at this stage are likely neural progenitors. Individual regions of interest (ROIs) were identified to investigate population-level activity patterns over time (Fig. 5A). Consistent with previous studies demonstrating spontaneous intracellular calcium fluctuations and coordinated calcium activity among ventricular zone progenitors (21), calcium signals at this stage were dominated by slow, sustained transients rather than rapid neuronal spiking. Individual ROI activity traces were binarized into active and inactive states, enabling quantification of the fraction of simultaneously active ROIs across the population over time (Fig. 5B).

**Figure 5.**
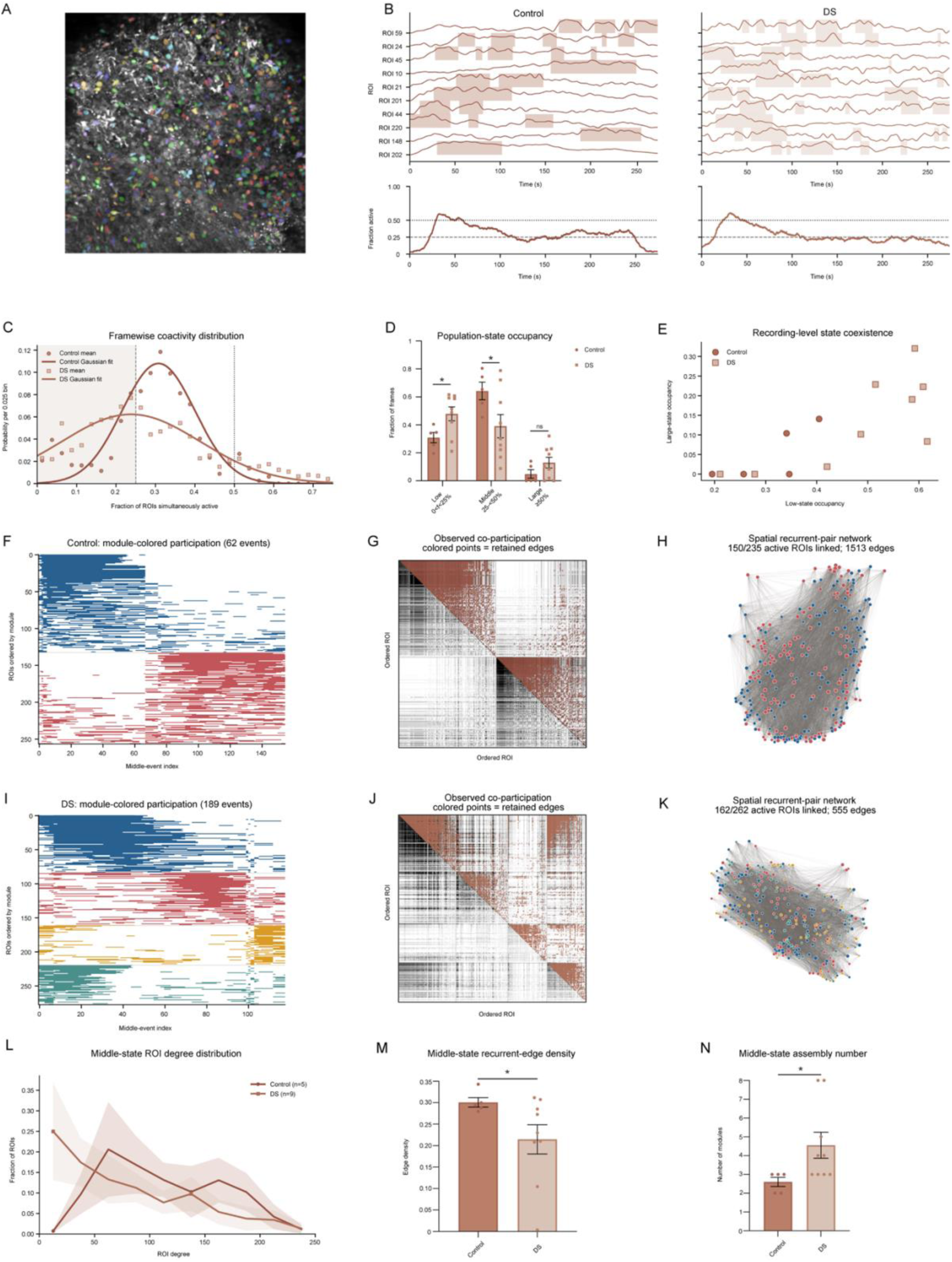
DS assembloids show altered population coactivity and reduced coordination within progenitor functional networks. (A) Representative calcium-imaging field with ROI overlays at day 30. (B) Representative smoothed ΔF/F traces and inferred active periods from control and DS assembloids. Lower traces show the fraction of all ROIs simultaneously active, f. (C) Assembloid-balanced distributions of f; curves are descriptive Gaussian fits. DS assembloids showed a broader distribution shifted towards lower coactivity values. (D) Fraction of recording time in low (0 < f < 0.25), middle (0.25 ≤ f < 0.50) and large (f ≥ 0.50) states. DS assembloids spent significantly more time in the low-coactivity state, while control ones spent significantly more time in the middle-coactivity state. (E) Low-versus large-state occupancy for individual assembloids. DS recordings displayed substantial occupancy of both low- and large-coactivity states. (F-K) Cellular organization of middle-coactivity events in representative control (F-H) and DS (I-K) assembloids. ROI participation across middle-coactivity events, ordered by consensus functional module (F, I). Event co-participation matrices; colored points denote ROI pairs with significant and consistent co-participation across middle-coactivity events (G, J). Spatial organization of the corresponding co-participating ROI pairs (H, K). Control assembloids contained denser patterns of ROI co-participation, whereas DS assembloids displayed a more dispersed organization. (L) Distribution of ROI degree, defined as the number of recurrent co-participating partners per ROI, during middle-coactivity events. (M) Density of consistently co-participating ROI pairs was lower in DS assembloids (control, 0.30 ± 0.01; DS, 0.21 ± 0.03). (N) DS assembloids contained more consensus functional modules (control, 2.60 ± 0.24; DS, 4.56 ± 0.69). Control, n=5; DS, n=9. Data are Mean ± S.E.M., ns: no significant changes observed, *P < 0.05-Welch’s t test. Each point on the graphs represents one assembloid. Co-participation relationships indicate statistical coordination across middle-coactivity events and do not imply direct anatomical connectivity. No inferential statistics were applied to the Gaussian fits in Fig. 5C or the representative examples in Fig. 5F-K. ROI, region of interest.

Analysis of population coactivity revealed distinct activity distributions between control and DS assembloids. Control assembloids exhibited a relatively narrow distribution concentrated around intermediate levels of population coactivity, whereas DS assembloids showed a broader distribution shifted towards lower coactivity values (Fig. 5C). To further characterize these differences, activity was classified into low-, middle-, and large-coactivity states. DS assembloids spent significantly more time in the low-coactivity state, while control ones spent significantly more time in the middle-coactivity state (Fig. 5D). Occupancy of the large-coactivity state was not significantly different between groups. Notably, increased low-state occupancy in DS assembloids did not simply reflect globally reduced activity. Instead, several DS recordings displayed substantial occupancy of both low- and large-coactivity states, indicating that these activity patterns could coexist within the same assembloid rather than representing distinct DS subgroups (Fig. 5E).

Previous work has shown that embryonic cortical progenitors do not function as isolated cells but instead form coordinated activity through gap junction-mediated coupling and functional networks (23, 24). We reasoned that the reduced occurrence of middle-coactivity events in DS assembloids might reflect an impaired ability to organize partially coordinated population activity. We therefore examined the cellular organization of these events by quantifying ROI participation across repeated middle-coactivity events and identifying ROI pairs that consistently co-participated across events. Representative raster plots showed repeated recruitment of specific ROI groups during middle-coactivity events in both control and DS assembloids (Fig. 5F, I). Recurrent co-participation relationships were visualized using co-participation matrices and spatial network reconstructions (Fig. 5G, H, J, K).

Qualitative analysis suggested that control assembloids contained denser and more recurrent patterns of ROI co-participation, whereas DS assembloids displayed a more dispersed organization. Consistent with this observation, the ROI degree distribution was shifted towards lower values in DS assembloids, indicating that individual ROIs participated in fewer recurrent co-participating pairs than in control assembloids (Fig. 5L). Likewise, the overall density of recurrent co-participation edges was significantly reduced in DS assembloids compared to controls (−30%; Fig. 5M), indicating that fewer ROI pairs repeatedly co-participated across events. Together, these findings suggest that coordinated recruitment within progenitor functional networks is weakened by *SCN1A* haploinsufficiency.

Despite this reduction in recurrent connectivity, DS assembloids contained significantly more functional modules than controls (77%; Fig. 5N). Thus, middle-coactivity patterns in DS organoids were distributed across a larger number of relatively discrete functional assemblies rather than being integrated into more densely interconnected networks. These findings are reminiscent of the functional organization described in developing cortical progenitors, where calcium activity emerges in coordinated network structures composed of recurrently interacting cell groups (24). Collectively, our data indicate that *SCN1A* haploinsufficiency does not simply reduce progenitor calcium activity but instead alters the organization of progenitor functional networks, resulting in increased low-state occupancy, reduced recurrent coupling among participating cells, and a more fragmented pattern of population activity.

To determine whether these network-level alterations reflected changes in the intrinsic electrophysiological properties of individual progenitors, we performed whole cell patch clamp recordings on assembloid sections at day 30 and injected biocytin into the patched cells, followed by immunostaining the sections for SOX2 and biocytin labelling to identify patched neural progenitors for quantifying electrophysiological parameters (Fig. S19A). Membrane capacitance, membrane resistance, and holding current were comparable between neural progenitors of control and DS assembloids (Fig. S19B-D), indicating that *SCN1A* haploinsufficiency does not primarily impair the intrinsic electrophysiological properties of ventricular zone progenitors. Rather, it disrupts the coordinated activity of progenitor populations, resulting in fragmented organization of calcium activity despite preserved single cell function.

## Discussion

DS has been majorly studied as a disorder of neuronal excitability arising from SCN1A-dependent dysfunction of inhibitory interneurons (3, 8, 12, 13, 34, 46). However, emerging evidence suggests that SCN1A may also function during early brain development (14–16). In the present study, we identify a previously underappreciated developmental role for SCN1A during human brain development. Using patient-derived forebrain assembloids arising from dorsal and ventral forebrain regions, we show that SCN1A is expressed within SOX2^+^ ventricular zone progenitors, where its haploinsufficiency is associated with reduced progenitor maintenance, accelerated cell cycle exit, premature neuronal differentiation, and increased excitatory neuron production. Furthermore, we demonstrate that *SCN1A* haploinsufficiency disrupts coordinated calcium dynamics across progenitor populations without altering the intrinsic electrophysiological properties of individual progenitor cells. Together, these findings suggest that SCN1A contributes to the organization of progenitor network activity during cortical development and that disruption of this function represents an early pathogenic mechanism in DS.

A major finding of this study is the identification of SCN1A expression at the apical surface of ventricular zone progenitors. Importantly, *SCN1A* haploinsufficiency resulted in progressive depletion of neural progenitors, reduced ventricular zone number and thickness, and impaired proliferative capacity. These defects occurred in the absence of increased apoptosis, indicating that altered progenitor behavior rather than enhanced cell death underlies the developmental abnormalities observed in DS assembloids. Our findings are further supported by recent observations in both mouse and human DS models reporting alterations in neurogenic niche function, neural stem cell maintenance, and progenitor cell cycle progression (16, 45).

The reduction in progenitor populations was accompanied by accelerated cell cycle exit and premature neurogenesis. DS assembloids exhibited fewer cycling progenitors, decreased progenitor proliferative fraction, increased neuronal output in pulse-chase experiments, and expansion of NeuN^+^ neurons. Notably, this expansion of neuronal population was driven predominantly by excitatory neurons, whereas inhibitory neuron numbers remained largely unchanged. These findings suggest that the loss of SCN1A function accelerates neurogenic output while simultaneously altering neuronal subtype composition. Although interneuron dysfunction is widely recognized as a central mechanism contributing to DS pathogenesis, our results indicate that developmental alterations in excitatory neuron production may represent a complementary pathogenic process. Consistent with this interpretation, increased excitatory neuron abundance was accompanied by enhanced neuronal network synchronization, increased delta power, stronger theta phase-locking, and spontaneous spiking activity at later developmental stages, which are hallmark events in epilepsy (47–53). Notably, no substantial differences in excitatory neuron composition were detected between control and DS samples in our single cell sequencing dataset. However, the increased abundance of excitatory neurons observed by immunofluorescence approaches suggests that subtype-specific changes may be masked within broad excitatory neuron classifications and will require higher resolution analyses to be fully resolved.

Single cell transcriptomic analyses provided further support for disrupted progenitor development. DS cycling radial glia, i.e., neural progenitors displayed advanced pseudotime positioning, increased developmental age, accumulation within S phase, and transcriptional signatures indicative of accelerated maturation. Particularly notable was the dysregulation of multiple connexin genes together with increased expression of calcium-responsive signaling genes. Gap junction-mediated communication is a critical component of the ventricular zone niche, coordinating calcium dynamics and progenitor behavior during corticogenesis (21, 24, 25, 55). Thus, the simultaneous alteration of connexin and calcium signaling pathways suggested that intercellular communication within progenitor populations may be disrupted by *SCN1A* haploinsufficiency.

Functional calcium imaging directly supported this prediction. Previous studies established that ventricular zone progenitors do not function as isolated cells but instead form interconnected networks of coordinated calcium activity. Owens and Kriegstein demonstrated spontaneous intracellular calcium fluctuations in cortical progenitors (21), while Bittman and colleagues showed that ventricular zone cells are organized into gap junction-coupled clusters whose coupling state changes dynamically during the cell cycle (23). Later, Malmersjö et al. demonstrated that neural progenitors form connexin 43-dependent small-world networks that promote proliferation and cortical growth (24). Our findings extend this framework by identifying SCN1A as a potential regulator of progenitor network organization. Rather than exhibiting a simple reduction in calcium activity, DS assembloids displayed increased low-coactivity states, decreased recurrent coupling, reduced network density, and increased segregation into discrete functional modules. Importantly, these abnormalities emerged at the population level despite preserved electrophysiological properties of individual progenitors, indicating that *SCN1A* haploinsufficiency primarily affects coordinated network behavior rather than cell autonomous electrical properties.

The convergence of transcriptomic and functional data raises the intriguing possibility that altered progenitor calcium dynamics contribute directly to the self-renewal defects observed in DS assembloids. Together, these findings suggest that *SCN1A* haploinsufficiency disrupts signaling mechanisms that normally coordinate progenitor maintenance and neurogenic timing.

Although interkinetic nuclear migration (IKNM) was not directly examined in the present study, our findings suggest a plausible mechanistic link between disrupted coordinated calcium activity and impaired progenitor self-renewal. Liu and colleagues demonstrated that gap junction-dependent calcium fluctuations regulate apically directed IKNM in ventricular zone progenitors and that disruption of connexin-mediated communication impairs nuclear translocation, alters mitotic positioning, and compromises maintenance of the progenitor pool (58). Given the altered expression of connexin genes observed in DS progenitors together with the fragmentation of progenitor functional networks identified here, it is reasonable to speculate that *SCN1A* haploinsufficiency similarly disrupts calcium-dependent regulation of IKNM in the neural progenitors. Such a mechanism would provide a direct link between altered progenitor communication and the downstream phenotypes observed in our model, including reduced self-renewal, premature cell cycle exit, ventricular zone depletion, and accelerated neuronal differentiation. This hypothesis offers a biologically plausible framework through which altered calcium network dynamics may influence corticogenesis in DS.

Based on these findings, we propose a working model (Fig. 6) in which SCN1A functions within apical ventricular zone progenitors to support coordinated calcium network activity. Loss of SCN1A disrupts gap junction-associated calcium activity, leading to fragmentation of progenitor networks and impaired coordination of progenitor behavior. We further speculate that these alterations interfere with calcium-dependent processes regulating IKNM and progenitor mitotic dynamics. Consequently, progenitor self-renewal is reduced, cell cycle exit is accelerated, and neuronal differentiation occurs prematurely. The resulting depletion of the progenitor pool drives excess excitatory neurogenesis and alters neural circuit assembly, ultimately promoting hyperexcitability and seizure susceptibility. Importantly, this model does not replace the established interneuron hypothesis of DS, but rather expands it by suggesting that developmental abnormalities in neural progenitors precede and contribute to later network dysfunction.

**Figure 6.**
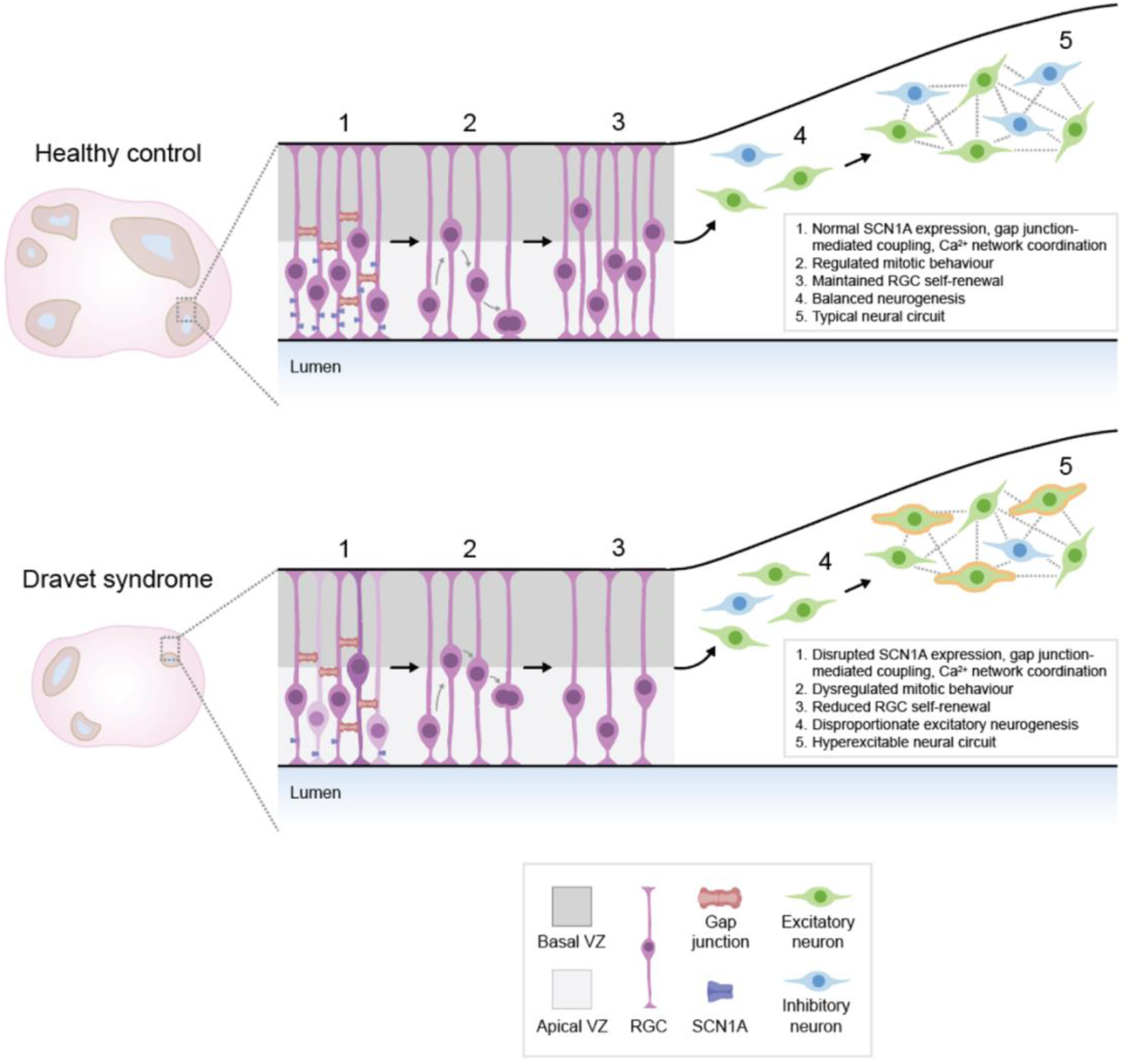
Proposed model linking *SCN1A* haploinsufficiency to disrupted progenitor network organization and altered neurogenesis in Dravet syndrome. Schematic illustrating the proposed developmental consequences of *SCN1A* haploinsufficiency. In healthy forebrain development (top), SCN1A supports gap junction-associated calcium-network coordination in ventricular zone radial glial cells, promoting regulated mitotic behavior, progenitor self-renewal, balanced neurogenesis, and normal neural circuit formation. In Dravet syndrome (bottom), reduced SCN1A expression disrupts progenitor calcium-network organization, leading to impaired progenitor maintenance, accelerated cell-cycle exit, premature excitatory neurogenesis, and hyperexcitable neural circuit formation.

Notably, certain limitations should be considered in our study. First, although assembloids recapitulate key aspects of human cortical development, they do not fully capture the cellular complexity and environmental influences present in vivo. Second, the proposed involvement of IKNM remains speculative and requires future BrdU-EdU incorporation studies to assess progenitor nuclear dynamics. Nonetheless, the convergence of cellular, transcriptomic, calcium imaging, and electrophysiological datasets supports a coherent developmental mechanism linking *SCN1A* haploinsufficiency to altered corticogenesis.

In summary, we identify a previously unrecognized role for SCN1A in regulating functional network organization within neural progenitor populations. Our findings suggest that *SCN1A* haploinsufficiency disrupts coordinated progenitor signaling, leading to impaired self-renewal, premature neurogenesis, excess excitatory neuron production, and hyperexcitable neural circuits. These results expand current models of DS pathogenesis beyond interneuron dysfunction and support the concept that early developmental abnormalities in ventricular zone progenitors contribute to disease onset. More broadly, our study identifies progenitor calcium network dynamics as a potential developmental mechanism linking *SCN1A* haploinsufficiency to altered corticogenesis and epilepsy, and highlights the developing ventricular zone as a potential therapeutic target in DS.

## Materials and methods

### PBMC isolation

All the protocols related to obtaining blood samples and subsequent iPSC/organoid generation were approved by Conjoint Health Research Ethics Board (#REB20-0251). Blood was drawn from a female patient (DS) and her healthy parents (mother; referred as control here and father; referred as control_3 here) by trained healthcare professionals at the Alberta Precision Labs in Foothills Medical Centre, Calgary. Peripheral blood mononuclear cells (PBMCs) were isolated from the blood samples using the Ficoll centrifugation protocol. Briefly, 3 ml blood was transferred to a sterile 15 ml conical tube and diluted 1:1 with sterile PBS. 4 ml Ficoll-Paque Plus solution (Sigma) was added to a fresh 15 ml tube and the diluted blood was carefully layered on top of it by keeping the tube in horizontal position.

Subsequently, the tube was placed back in a vertical position. The tube was centrifuged for 30 minutes at 1,000 xg at room temperature without brake. The top most aqueous phase (plasma) was carefully removed. The buffy coat (white layer between the Ficoll solution border and aqueous phase) was collected and transferred to a sterile tube. The buffy coat was washed with PBS by centrifugation for seven minutes at 540 xg at room temperature with the highest brake. The pellet obtained was resuspended in PBS and cell counting was performed using trypan blue on the cell counter (Countess, Invitrogen). The cell pellet was further diluted with PBS and centrifuged for five minutes at 300 xg at room temperature. The final pellet containing PBMCs was resuspended in 1 ml freezing solution (90% fetal bovine serum, 10% DMSO) and transferred into cryovials that are stored in −80°C for 24 hours before moving to liquid nitrogen storage tank.

### iPSC reprogramming and validation

PBMCs obtained from patient (DS) and her healthy parents (control and control_3) were reprogrammed into iPSCs using CytoTune-iPS2.0 Sendai Reprogramming Kit (Thermo Fisher Scientific). The iPSCs were validated by immunostaining for pluripotency marker antibodies (OCT3/4, SOX2, SSEA-4, TRA-1-60). Briefly, the cells were fixed in 4% paraformaldehyde (PFA) for 15 minutes at room temperature, followed by washing with DPBS, permeabilization (DPBS/0.1% Triton X-100) for 15 minutes and blocking (DPBS/10% fetal bovine serum) for one hour at room temperature. Later, the cells were incubated in primary antibodies overnight at 4°C, followed by washing and secondary antibody incubation overnight at 4°C. Finally, the immunostained cells were washed and counterstained with Hoechst (Thermo Fisher Scientific). Using published protocol (59), simple and cystic EB aggregates were generated and allowed to differentiate in culture. Immunostaining for in vitro differentiation marker antibodies (AFP, βIII-tubulin, and α-SMA) was performed as described above. Brightfield and fluorescent imaging was performed using Zeiss Axio Scope A1. The primary antibodies (and their working concentrations) used in this study are listed in Table S1. Secondary antibodies were used at 1:200 concentration (Life Technologies).

iPSCs generated from an unrelated female patient (DD1C; referred as DS_2 here) were kindly shared by Jens Schuster (42) and iPSCs generated from an unrelated healthy female individual (ACS-1023; referred as control_2 here) were obtained commercially (Cedarlane Labs).

### iPSC maintenance

iPSCs were plated on matrigel-coated plates (Matrigel Matrix hESC-Qualified Mouse, VWR) and maintained in mTeSR Plus media containing 0.5X Penicillin-Streptomycin (Life Technologies). Briefly, thawed cells were gently added to a 15 ml tube containing 9 ml pre-warmed mTeSR Plus media and centrifuged at 200 xg for 5 minutes at room temperature. Cell pellet was resuspended in 1 ml mTeSR Plus media and cell counting was performed. 300,000 cells were plated on 60 mm tissue culture plate (VWR) in a final volume of 3 ml media containing 10 μM Y-27632. The plate was gently placed into the 37°C/5% CO_2_ incubator. After 24 hours, cells were observed under the microscope and the spent media was replaced with fresh pre-warmed mTeSR Plus media. Cells were maintained until 70% confluency and were passaged using accutase cell dissociation reagent (Thermo Fisher Scientific).

### Organoid and assembloid generation and fusion

Forebrain organoids (hCO and hMGEO) were generated from the patient and healthy control iPSCs by adapting the protocol from Xiang et al. (31). Briefly, iPSCs were dissociated using accutase, EBs were cultured in 96-well plates (U-bottom, ultra-low attachment, Corning), hCO/hMGEO were generated at day 10 and fused at day 18 to form assembloids which were maintained in ND media in 6-well plates (clear bottom, ultra-low attachment, Corning) on an orbital shaker (80 rpm) in 37°C/5% CO_2_ incubator. The media and reagents used in the culture are mentioned in the protocol.

### Immunofluorescence, EdU labelling and AAV labelling

Assembloids were fixed in 4% PFA overnight at 4°C, followed by washing with PBS and soaking in 20% sucrose at 4°C until they sink to the bottom of the tube. Later, the assembloids were embedded in OCT and stored at −80°C, followed by obtaining 12 µm thick cryosections. To perform immunostaining (for PAX6, TTF-1, Calbindin, SCN1A, SOX2, ZO-1, Ki67, NeuN, vGLUT1, GAD67, CC3 and GABA) on cryosections, antigen retrival was carried out with citrate buffer (pH 6.0) for 15 minutes at 95°C, followed by permeabilization with PBS/1% Triton X-100 for 30 min at room temperature and blocking in 5% normal donkey serum for 1 hour at room temperature. Later, the sections were incubated in primary antibody overnight at 4°C, followed by washing and secondary antibody incubation for 2 hours at room temperature. Finally, the immunostained sections were washed, counterstained with Hoechst and mounted for imaging. The primary antibodies (and their working concentrations) used in this study are listed in Table S1. Secondary antibodies were used at 1:200 concentration (Life Technologies). For experiments involving EdU co-staining, the sections were incubated with EdU Click-iT reaction mix (prepared fresh as per manufacturer’s instructions; Invitrogen) for 30 minutes in the dark after secondary antibody incubation and washing steps. For pAAV-CAG-tdTomato (59462, Addgene) viral labelling, day 15 hMGEOs were incubated in 250 µl ND media/5 µl AAV (1.9 × 10^13^ gc/ml) in a ultra-low attachment 96-well plate and the plate was gently placed into the 37°C/5% CO_2_ incubator. After three days (day 18), the AAV labelled hMGEOs were washed several times with fresh ND media. This was followed by adding unlabelled hCO to the hMGEO-containing wells in 96-well plate and placing the plate back in the incubator for three days of fusion. At day 21, i.e., 3 daf, the assembloids were transferred to an ultra-low attachment 6-well plate containing fresh ND media and placed on an orbital shaker (80 rpm) in the 37°C/5% CO_2_ incubator. The assembloids were fixed for cryosectioning at day 28 (10 daf) and the sections (containing endogenous tdTomato signal) were used for immunofluorescence as described above.

### Imaging and quantification

Brightfield images of whole EBs and assembloids were obtained using inverted microscope (Zeiss Primovert). The immunostained cryosections were imaged using LSM 880/LSM 900 confocal microscopes (Zeiss). Three sections per assembloid were imaged and each section was imaged in four quadrants. Immunostaining for TTF-1 antibody was performed to distinguish quadrants as hCO (TTF-1^-^) and hMGEO (TTF-1^+^). Cell counting was performed with ZEN software (Zeiss) and quantifications were performed separately on hCO and hMGEO regions. Size measurements were performed using ZEN software.

### Electrophysiological analysis

Electrophysiological recordings were obtained from assembloids at day 90. For each recording session, an assembloid was removed from ND media, transferred to a custom-made recording chamber, and partially submerged in a drop of 1.5% (w/v) low-melting-point agarose to minimize movement. During recording, assembloids were continuously perfused with mildly elevated-potassium (8.5 mM K^+^) artificial cerebrospinal fluid (ACSF) to enhance network activity (60). A 64-channel silicon probe (Cambridge Neurotech ASSY-77-E-1, Cambridge, UK) was inserted into the assembloid using a motorized micromanipulator (Luigs and Neumann, Ratingen, Germany). A silver wire placed adjacent to the assembloid served as the grounding electrode. Neural signals were amplified with an RHD2132 headstage (Intan Technologies, Los Angeles, CA, USA) and digitized at 30 kHz using an Open Ephys USB acquisition board (Open Ephys, Atlanta, GA, USA). Hardware filters were set to 1–7,000 Hz, and raw data were acquired using the Open Ephys GUI v0.5.5.3. Each assembloid was recorded for three 10-minute sessions at spatially separated locations.

Data analyses were performed in Python 3.10.16 using SpikeInterface (61) and custom analysis scripts. For consistency, analyses were restricted to the first 10 minutes of each recording. The acquisition channel map was aligned to the geometry of the Cambridge Neurotech ASSY-77-E-1 probe. Global artifacts were detected using a 500-3000 Hz filtered copy of the raw signal. Events exceeding 25 times the channel-wise robust standard deviation on at least 10 channels were classified as artifacts, expanded by 5 ms on either side, and excluded from downstream analyses without interpolation.

For LFP power analysis, signals were band-pass filtered from 1-100 Hz, notch filtered at 60 Hz, globally common-average referenced, and downsampled to 1000 Hz. Power spectral density was calculated from artifact-free Welch windows in the delta (1-4 Hz), theta (4-8 Hz), alpha (8-12 Hz), beta (12-30 Hz), low-gamma (30-55 Hz), and high-gamma (65-100 Hz) bands (Fig S20). Pairwise magnitude-squared coherence was calculated in the high-gamma band (65–100 Hz) without common-average referencing, using 1 s Hann windows with 50% overlap. Inter-electrode distances were calculated from probe geometry, and coherence was summarized in 100 µm distance bins (Fig. S14A).

MUA was isolated using a 300-6000 Hz band-pass filter and global median referencing. Negative peaks were detected at five times the artifact-free noise estimate (Fig S20). Spikes occurring within 1.5 ms across neighbouring electrodes were merged to reduce duplicate detections. Network bursts were defined as sequences of at least three spikes separated by <100 ms and were retained when they lasted >50 ms and recruited >10% of valid channels. Network firing rate, burst metrics, inter-burst LFP power, burst-triggered LFP responses, and theta phase locking were then quantified. Theta-MUA phase coupling was expressed as the mean resultant length of spike-associated theta phases within continuous artifact-free segments.

For single-unit analysis, recordings were assigned the probe geometry and, for consistency, analyses were restricted to the first 10 minutes of each recording. Signals were band-pass filtered from 300-6000 Hz, notch filtered at 60 Hz, globally common-average referenced, and sorted with GPU-accelerated Kilosort4 (62). Kilosort4 was run with Th_learned = 9, Th_universal = 10, min_template_size = 25, acg_threshold = 0.2, n_templates = 6, and n_pcs = 6. For recordings with insufficient waveform clips for this initialization, the number of templates, and, when necessary, principal components, were reduced only to the number supported by the available clips; recordings with fewer than two usable clips were retained as zero-unit recordings for recording-level analyses. Following sorting, spikes within any 5-ms window containing activity from at least 10 distinct units were removed as putative synchronous artifacts, and units with fewer than 10 remaining spikes were discarded. Units were iteratively hard-merged when template similarity exceeded 0.90 and the Euclidean distance between monopolar-triangulation location estimates was <40 µm; connected merge groups were resolved iteratively until no further pairs met both criteria. Up to 500 randomly selected spikes per unit (random seed = 42) were used for waveform and template estimation. Final quality curation retained units with a signal-to-noise ratio >4 and an interspike-interval (ISI) violation ratio <0.1. Unit-level measures included spike count, mean firing rate, median absolute extracellular amplitude, ISI coefficient of variation (CV) and CV2, within-unit burst fraction (the proportion of consecutive ISIs <10 ms), template trough-to-peak duration, and template spatial spread, defined as the number of channels with a peak-to-peak template amplitude of at least 20% of the maximum channel amplitude.

Statistical analyses were performed in Python 3.10.16. Group comparisons were modelled with group as the fixed effect and control assembloids as the reference group. Model coefficients therefore represent the estimated difference between DS and control aseembloids, reported as DS minus control. Recording-level LFP, coherence, MUA, and spike-sorting yield/population measures were analyzed using mixed-effects models with assembloid identity included as a random intercept to account for repeated recordings from the same assembloid. Absolute power measures were analyzed on the log10 scale, whereas relative power and coherence measures were analyzed on the logit scale. Single-unit metrics were analyzed using mixed-effects models with assembloid identity as a random intercept and recording identity as an additional variance component, accounting for units nested within recordings and assembloids. Spike counts, firing rates, ISI CV, ISI CV2 (63), and template spatial spread were log1p-transformed; extracellular amplitude and template trough-to-peak duration were log10-transformed; and active-channel and within-unit burst fractions were logit-transformed. When mixed-effects models failed to converge or produced singular fits, transformed data were aggregated to assembloid-level means and compared using Welch’s t-test. Multiple comparisons were controlled within pre-specified frequency-band or related metric families using the Benjamini-Hochberg false-discovery-rate (FDR) procedure. Spatial coherence decay was assessed using 100-µm distance bins by modelling logit-transformed coherence as a function of interelectrode distance, group, and their interaction; when the spatial mixed model was unstable, one coherence-distance slope was estimated per assembloid, and slopes were compared between groups using Welch’s t-test. Statistical significance was evaluated at FDR-adjusted Q < 0.05.

### Cross-correlation analysis

Spike-time cross-correlation histograms (CCHs) were calculated using Elephant 1.2.1 (64) and Neo 0.14.5. Spike trains from each 600 s recording were discretized into 5 ms binary bins, and pairwise CCHs were calculated over lags from −100 to +100 ms and normalized as cross-correlation coefficients (65). No smoothing or border correction was applied. For each unit pair, the maximum and minimum CCH coefficients within the lag window were retained.

For descriptive thresholding, CCH extrema were pooled across all 537-unit pairs without reference to group labels. High-positive relationships were defined as pairwise maxima at or above the pooled 75^th^ percentile (0.0142663), and high-negative relationships as pairwise minima at or below the pooled 25^th^ percentile (−0.00222145). These thresholds were used for visualization and calculation of the high-positive CCH fraction and were not statistical significance thresholds.

For each recording containing at least four units, the high-positive CCH fraction was calculated as the proportion of unit pairs meeting the pooled 75^th^-percentile criterion. Mean positive CCH peak and mean negative CCH trough were calculated for each recording as the arithmetic means of the pairwise CCH maxima and minima, respectively. Recording-level values for these three measures were averaged within each assembloid, yielding 9 control and 5 DS assembloids. Groups were compared using unpaired two-sided Welch’s t-tests.

No surrogate-based pairwise significance test was performed. Accordingly, CCH relationships were interpreted as descriptive spike-time associations rather than validated synaptic connections, and CCH sign was not used to assign excitatory or inhibitory neuronal identity.

Electrophysiological recordings were performed on day 150 assembloids by placing them on an upright Olympus BX51 microscope, visualized using a 5X Olympus N-Achroplan objective, and continuously perfusing with carbogen-bubbled fluid ACSF. All recordings were performed at 36 ± 1 °C. A glass micropipette (3–8 MΩ) filled with ACSF was lowered into the assembloid. Extracellular field potentials were acquired in current-clamp mode. Recordings were low-pass filtered at 1 kHz and high-pass filtered at 0.1 Hz with a Bessel filter, and digitized at 10 kHz (Multiclamp 700B amplifier, Digidata 1440A digitizer, Axon Instruments), and stored using pClamp software (Axon Instruments). Spontaneous events were detected offline in Clampfit (Axon Instruments). The threshold for detection was set at three times the noise (peak-to-peak amplitude). Peak amplitude (Mean ± S.D.) and event frequency were calculated for each recording that contained detectable spikes. Each assembloid was recorded at multiple locations to account for spatial heterogeneity. Spike frequency and amplitude were quantified from the most active recording obtained in each assembloid.

### Single cell suspension preparation

Single cell dissociation of pools of 6-8 assembloids per sample at day 90 was performed using the Worthington Papain Dissociation kit manufacturer’s protocol at 37°C. Briefly, assembloids were transferred to a 60 mm dish, Papain and DNase solution were added to the dish, and assembloids were minced into small pieces using a razor blade. The dish was transferred to the orbital shaker inside a humidified tissue culture incubator for 30 minutes. The minced pieces were pipetted up-and-down using a 10 ml pipette. The cell suspension (excluding the debris) was transferred to a 15 ml tube containing Earle’s medium plus inhibitor solution. Cells were centrifuged for 5 minutes and the pellet was resuspended in 1x PBS/0.5% BSA. The resuspended cells were passed through 40 μm nylon filter. Cell counting was performed and viable cells were FACS-sorted after treating with the viability dye (eBioscience Fixable Viability Dye eFluor 780).

### scRNA-seq library construction and sequencing

Cells for each sample were processed with Chromium Next GEM Single Cell 3’ v3.1 Chemistry (10x Genomics) as per the manufacturer’s protocol. In brief, single cells in 1x PBS/0.5% BSA were partitioned into Gel bead-in-Emulsions (GEMs) using GemCode Technology. DynaBeads MyOne Silane magnetic beads (Invitrogen) were used to remove leftover biochemical reagents, then cDNA was amplified by PCR over 12 cycles. Read 1 primer sequences were added to cDNA during GEM incubation. P5 primers, P7 primers, i7 sample index, and Read 2 primer sequences were added during library construction. Library quality and fragment size distribution were assessed using the TapeStation D1000 ScreenTape assay (Agilent). Sequencing was performed over multiple rounds using the NovaSeq 6000 platform (Illumina) at the University of Calgary Centre for Health Genomics and Informatics (CHGI). Approximately 15,000 to 60,000 cells were recovered per sample, with a targeted sequencing depth of 20,000 reads per cell. Sequencing reads were aligned to the pre-built human reference genome (GRCh38) and aggregated into a single dataset using the CellRanger (version 8.0.1) pipeline (66). The aggregated dataset contained 202,854 cells with 15,751 post-normalization reads per cell.

### scRNA-seq data analysis

#### Pre-processing, dimensional reduction, clustering, and visualization

The aggregated feature-barcode matrix was imported into Seurat (version 5.3.0) (67) for downstream normalization, variable feature identification, data scaling, integration, clustering, dimensionality reduction, and visualization with default parameters. Quality control metrics were visually inspected, and poor quality cells and suspected multiplets (>7000 detected genes, >22,000 detected UMIs, and/or >10% mitochondrial reads) were filtered out. Individual samples were integrated using Harmony (version 1.2.3) (68), with clustering and dimensionality reduction subsequently performed using the top 50 principal components. Cluster identities were manually annotated by querying previously characterized markers from developing human brain and brain organoid scRNA-seq datasets (29, 69–72), and corroborated by examining DEGs in each cluster. To focus analysis on neural cell type populations found in the brain, putative immune cells (*TYROBP+*, *C1QA+*), retinal cells (*TPH1+*, *SMIM36+*), mesenchymal cells (*COL1A1+*, *COL3A1+*), and melanocytes (*MITF+*, *DCT+*) were removed by iteratively subsetting and reprocessing the dataset through this standard workflow.

#### Identification of DEGs, Gene Ontology (GO) enrichment, and cell cycle phase

Cluster markers conserved across samples and DEGs between samples (expressed in at least 10% of cells in the given cluster or group of interest and with a minimum log2 fold change threshold of 0.25) were identified via the Wilcoxon rank-sum test implemented in FindAllMarkers and FindMarkers. Gene Ontology (GO) term enrichment analysis was performed with DEenrichRPlot, a Seurat wrapper around Enrichr (73), using a maximum of 300 input genes and the Biological Process 2025 annotations. Cell cycle scores were determined using CellCycleScoring, which assigned each cell as being in G1, S, or G2M phased based on the expression of a curated list of cell cycle markers (74).

#### RNA velocity

Developmental trajectories within the excitatory lineage were examined by RNA velocity analysis. CellRanger count-generated BAMs for each sample were realigned with velocyto (version 0.17.17) (75) with the run10x command and human (GRCh38) annotations. The output loom files containing spliced and unspliced counts for each control and DS sample were combined and imported into scVelo (version 0.2.5) (76) along with pre-computed metadata (i.e., sample, cluster identity, and UMAP coordinates) from an integrated Seurat object which was subsetted and processed as above to contain only cycling RGCs, RGCs, IPCs, and excitatory neurons. The standard scVelo workflow was applied with default parameters, and RNA velocities were estimated using the dynamical model and visualized as streamlines. Calculated velocity pseudotime values were exported from the AnnData object metadata and visualized as kernel density plots using ggplot2 (version 3.5.2).

#### Developmental age prediction

The developmental age of cell types from control and DS assembloids was predicted using a published cell type-agnostic model (54). Normalized counts were pseudobulked by individual sample replicate (2 replicates for control, 2 replicates for control_3, 2 replicates for DS, 1 replicate for DS_2) and cell type, and genes present in the model but not detected in the query pseudobulked dataset were zero padded.

Sequencing raw data are available at NCBI’s Gene Expression Omnibus (GEO accession: GSE341547) and are currently private until published.

### Calcium imaging and analysis

Assembloids at day 30 were incubated in ND media containing Fluo-4 AM (Thermo Fisher Scientific) and Pluronic F-127 (Thermo Fisher Scientific) for one hour in 37°C/5% CO_2_ incubator, followed by washing with PBS and ND media. The assembloids were later incubated in freshly prepared ACSF at 37°C and live calcium imaging was performed with time series recordings (3000 cycles, 11 frames/s) in LSM 880 confocal microscope Airyscan FAST mode (Zeiss).

#### Calcium activity-state analysis

ROI-wise ΔF/F traces from 5 control and 9 DS assembloids were analyzed (one recording per assembloid; 3,000 frames at 10.94 Hz). Traces were Gaussian-smoothed (σ = 2 s) and converted to binary active states using Schmitt-trigger thresholding (77–79). The upper and lower thresholds were set to the 80^th^ and 60^th^ percentiles of each ROI trace, respectively. Inactive gaps ≤0.5 s were filled and active periods < 3 s were removed.

Population coactivity was defined as the fraction of active ROIs at each time point,

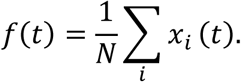

Each frame was classified as low (0 < *f* < 0.25), middle (0.25 ≤ *f* < 0.50), or large (*f* ≥ 0.50) coactivity. Frames with *f* = 0 were unclassified. State occupancy was calculated as the fraction of all recording frames in each state. Robustness to the activity-state parameters, including the thresholds used to define ROI activity and the population coactivity-state boundaries, was assessed in Fig. S21. For Fig. 5C, values with *f* = 0 or *f* = 1 were excluded, binned at 0.025 intervals, and normalized within each organoid before group averaging. Gaussian fits to the group-mean distributions were used for visualization only.

#### Middle-coactivity events and ROI co-participation

Middle-coactivity events were identified from local maxima of *f*(*t*) within middle periods (0.25 ≤ *f* < 0.5). Local maxima were required to be separated by at least three frames, and one maximum was retained from each contiguous middle-coactivity period. ROI participation was represented by a binary ROI × event matrix *X*, and pairwise co-participation counts were calculated as *XX*^T^.

Following previous coactivity-based functional-network analyses in calcium imaging data (80, 81), recurrent ROI co-participation was assessed against surrogate activity patterns. For each assembloid, 1,000 surrogate event matrices were generated using the Curveball algorithm (82), preserving both ROI participation frequency and event size. ROI pairs with an empirical one-sided P < 0.01 were retained as functional edges, consistent with thresholds previously used to infer functional connections from calcium-imaging data (83). These edges represent statistically recurrent co-participation and do not imply anatomical connectivity.

#### Functional modules and co-participation measures

Retained ROI pairs were represented as unweighted, undirected graphs. Functional modules were identified using the Leiden algorithm (84) with the Reichardt–Bornholdt configuration-model objective (85) and a resolution parameter of *γ* = 1. Isolated ROIs were excluded from community detection and from the module count.

For Fig. 5L, ROI degree was defined as the number of retained functional edges per ROI. Degree distributions were calculated across all ROIs within each assembloid, binned in intervals of 25 edges, and expressed as the fraction of ROIs in each bin before group averaging. Edge density (Fig. 6m) was *E*/(*^N^*), where *E* is the number of retained pairs and *N* is the total number of ROIs. Module number (Fig. 5N) was defined as the number of Leiden communities among non-isolated ROIs.

### Patch clamp

300 µm vibratome sections of assembloids were placed in a patch-clamp electrophysiology chamber at day 30 and constantly perfused with an extracellular solution containing (in mM): NaCl (50), CaCl_2_ (1), MgCl_2_ (1), CsCl (1), CdCl_2_ (0.3), tetraethylammonium-Cl (100), 4-aminopyridine (1), HEPES (10) and glucose (10) at pH 7.4 (with CsOH). Cells were whole-cell patch-clamped with a holding potential (V_H_) of −60 mV. The pipette solution contained (in mM): Cs-methanesulfonate (120), NaCL (10), MgCl_2_ (1), HEPES (10), Mg-ATP (2), Na-GTP (0.2) and biotin 4-amidobenzoic acid sodium salt (12.9) at pH 7.2 (CsOH). Biocytin (Invitrogen) was injected into the cells and they were visualized on a Nikon BX51 microscope. Data were collected at 10 kHz and lowpass Bessel filtered at 2 kHz using an Axopatch 700B and Digidata 1550 (Molecular Devices). The membrane test feature of pClamp 11 (Molecular Devices) was used to determine the basic membrane properties: holding current, membrane capacitance (C_m_) and membrane resistance (R_m_). The free-floating patched sections were fixed in 4% PFA, followed by immunostaining for SOX2 and co-labeling for Biocytin using Alexa Fluor 488-conjugated Streptavidin (1:200, Jackson ImmunoResearch Laboratories Inc.). The stained sections were mounted on slides and imaged using LSM 880 confocal microscope (Zeiss). SOX2^+^/Biocytin^+^ cells were identified as neural progenitors that were patched.

### Statistical analysis

GraphPad Prism software was used to perform statistical analysis unless stated otherwise. Data are represented as Mean ± S.E.M. and Mean ±S.D. Statistical tests used in the experiments are mentioned in the corresponding figure legends.

## Author contributions

D.D. and D.M.K. were responsible for conception and design of the research. D.D. performed the experiments, analyzed the data and wrote the initial draft of the manuscript. H.F. performed transcriptomic data analysis and contributed to generating the working model. S.Z. and C.Z. performed electrophysiological experiments and analyzed the data. S.J.G. provided electrophysiological analysis support. T.S. analyzed the calcium imaging data and contributed to electrophysiological data analysis. N.R., N.C. and J.B. contributed to running scRNA-seq reactions and contributed to data analysis. M.G. and A.K. contributed to immunostaining experiments. D.N. and R.T. contributed to patch clamp experiment and data analysis. L.R. and D.R. provided early iPSC/organoid culture facility support, shared reagents and provided troubleshooting insights. J.J.L. collaborated by providing access to blood samples and shared expert suggestions on epilepsy research.

## Acknowledgments

We would like to thank the Kurrasch lab members for their input during this study. We also thank Christine Cheng (NTU Singapore) for kindly sharing PBMC isolation protocol. We thank Kingsley Ibhazehibo for help with PBMC isolation from DS and control blood samples. We also thank Shiying Liu and Centre for Genome Engineering (UCalgary) for providing iPSC reprogramming and validation services. We thank Elodie Labit and Jessica Joon for providing FACS-sorting services. We further thank Natasha Klenin and Rachel Oskin for help with cryosectioning. We thank Jens Schuster (Uppsala University) for generously sharing DD1C iPSC line. We also thank Sisu Han for providing guidance on calcium imaging. We thank Pia Svendsen (ACHRI) for training on confocal microscopes. We are grateful to Alicia Vandenbrink and Kristy Gough for all the administrative support. Finally, we appreciate Linden (DS patient) and her family for their continued interest in our research program.

