## Supplementary information for "*SCN1A* haploinsufficiency disrupts neural progenitor networks and accelerates neurogenesis in human Dravet syndrome forebrain assembloids"

[1]Department of Medical Genetics, University of Calgary, Calgary, Alberta, Canada
[2]Department of Comparative Biology and Experimental Medicine, University of Calgary, Calgary, AB, Canada
[3]Department of Pediatrics, University of Calgary, Calgary, Alberta, Canada
[4]Department of Cell Biology and Anatomy, University of Calgary, Calgary, Alberta, Canada
[5]Cambridge NeuroTech
[6]Department of Biochemistry and Molecular Biology, University of Calgary, Calgary, Alberta, Canada
[7]Alberta Children's Hospital Research Institute, University of Calgary, Calgary, AB Canada
[8]Hotchkiss Brain Institute, University of Calgary, Calgary, AB, Canada

### Supplementary figure legends

**Figure S1. Validation of pluripotency and trilineage differentiation potential of iPSC lines.** (A) Immunofluorescence-based pluripotency characterization of control and DS iPSC lines at passage 6. Both control and DS iPSC colonies express the canonical pluripotency markers OCT3/4 (green), SOX2 (green), SSEA-4 (red), and TRA-1-60 (green). Nuclei were counterstained with Hoechst (blue). 3-6 clones were tested for each iPSC line. (B) Assessment of trilineage differentiation potential using EB-based differentiation assays. Brightfield images of simple and cystic EB aggregates are shown in the top panel. Representative images show the immunofluorescence-based expression of α-fetoprotein (AFP; endoderm), βIII-tubulin (ectoderm), and α-smooth muscle actin (α-SMA; mesoderm) in differentiated control and DS EBs. These findings confirm the capacity of both control and DS iPSC lines to generate derivatives of all three germ layers. 3-4 clones were tested for each iPSC line. DS, Dravet syndrome; iPSCs, induced pluripotent stem cells; EBs, embryonic bodies.

**Figure S2. Schematic overview of forebrain assembloid generation protocol.** (A) Timeline illustrating the generation of hCOs, hMGEOs, and forebrain assembloids from iPSCs. iPSCs were maintained in mTeSR Plus medium prior to EB formation (day 0). Neural induction was initiated using neural induction medium from day 0 to day 10, followed by dorsal and ventral forebrain patterning using ND medium without vitamin A (hCOs) or MGE patterning medium (hMGEOs) from day 10 to day 18. Region-specific organoids were subsequently matured and fused to generate forebrain assembloids. Following fusion, assembloids were maintained in ND medium for long-term culture and maturation. Time points corresponding to iPSC expansion, EB formation, organoid specification, neural maturation and fusion, and long-term assembloid culture are indicated. hCO, human cortical organoids; hMGEO, human median ganglionic eminence organoids; ND, neural differentiation.

**Figure S3. DS forebrain assembloids exhibit regional patterning and interneuron migration.** (A) Validation of dorsal-ventral regional specification in DS assembloids at 3 daf (day 21). Immunostained sections demonstrate confinement of the dorsal forebrain progenitor marker PAX6 (red) to the hCO region and the ventral forebrain marker TTF-1 (NKX2.1; green) to the hMGEO region. DS, n=3. (B) Validation of interneuron migration in DS assembloids using AAV-CAG-tdTomato-labelled hMGEOs. At 10 daf (day 28), tdTomato$^+$ cells (red) originating from the TTF-1$^+$ (green) hMGEO region are observed migrating into the adjacent hCO region. DS, n=3. daf, days after fusion.

**Figure S4. Independent validation of reduced growth in DS forebrain assembloids using a second patient-derived iPSC line.** (A) Representative brightfield images of control_2 and DS_2 EBs (day 10) and assembloids (day 60), illustrating reduced growth of DS cultures. (B, C) Quantification of EB and assembloid size during development. DS cultures exhibit significantly reduced size relative to controls at day 10 (B) and day 60 (C). Control_2 and DS_2, n=3 each genotype for day 10; n=3 each genotype for day 60. Data are Mean ± S.D., *$P < 0.05$, **$P < 0.01$- Welch's t test. Each point on the graphs represents one assembloid.

**Figure S5. *SCN1A* is expressed in fetal human radial glial progenitor populations.** Dot plot showing expression of *SCN1A* together with the neural progenitor markers SOX2 and HES1 across radial glial cell populations in a published human fetal brain single cell transcriptomic dataset. Cell populations include early radial glia (RG-early), outer radial glia (oRG), ventricular radial glia (vRG), transitional radial glia (tRG), dividing radial glia (RG-div1 and RG-div2), and medial ganglionic eminence radial glia (MGE-RG1 and MGE-RG2). Dot size represents the proportion of cells expressing each gene, while color intensity indicates average normalized gene expression. *SCN1A* expression is detectable across multiple radial glial progenitor populations, supporting the presence of *SCN1A* in neural progenitors during human fetal brain development.

**Figure S6. SCN1A is expressed in both excitatory and inhibitory neurons in mature forebrain assembloids.** Representative immunofluorescence images of day 90 control assembloids showing SCN1A expression (magenta) in both excitatory and inhibitory neuronal populations. Co-immunostaining with the excitatory neuron marker vGLUT1 (green; left panel) and the inhibitory neuron marker GAD67 (green; right panel) demonstrates SCN1A expression in multiple neuronal subtypes within mature assembloids. Arrowheads indicate representative vGLUT1$^+$/SCN1A$^+$ and GAD67$^+$/SCN1A$^+$ cells. Control, n=3.

**Figure S7. DS assembloids exhibit reduced size and disrupted ventricular zone architecture.** Representative whole-section images of day 30 assembloids stained with nuclear marker Hoechst (magenta) to visualize overall tissue organization and ventricular zone-like structures. Compared with control and control_2 assembloids, both DS and DS_2 assembloids display a reduced overall size and fewer ventricular zone-like rosettes. Similar architectural abnormalities were observed in two independent DS patient-derived lines, supporting a reproducible effect of *SCN1A* haploinsufficiency on early forebrain development. Control, n=3; Control_2, n=3; DS, n=3; DS_2, n=3.

**Figure S8. Reduced neural progenitor populations in an independent patient-derived assembloid.** (A) Representative immunofluorescence images of day 60 assembloids generated from control_2 and DS_2, stained for the neural progenitor marker SOX2 (green) and Hoechst (magenta). DS_2 assembloids exhibit a marked reduction in $SOX2^+$ progenitor cells compared with control_2 assembloids. (B) Quantification of $SOX2^+$ cells in the hCO and hMGEO regions at day 60. DS_2 assembloids display a significant reduction in the percentage of $SOX2^+$ neural progenitors (out of total $Hoechst^+$ cells) in both forebrain compartments relative to control_2 assembloids, independently validating the progenitor depletion phenotype observed in the primary DS line. Control_2, n=4 for hCO and hMGEO; DS_2, n=4 for hCO and hMGEO. Data are Mean ± S.D., ns: no significant changes observed, *P < 0.05, ***P < 0.001- Welch's t test. Each point on the graph represents one assembloid.

**Figure S9. *SCN1A* haploinsufficiency disrupts ventricular zone maintenance during forebrain assembloid development.** (A) Representative immunofluorescence images of control and DS assembloids stained for SOX2 (green) at day 30, day 60, and day 90. Dashed outlines indicate ventricular zone-like structures. DS assembloids exhibit fewer and progressively smaller $SOX2^+$ ventricular zones compared with control assembloids. (B–D) Quantification of the number of ventricular zones per assembloid section at day 30 (B), day 60 (C), and day 90 (D). DS assembloids display a significant reduction in ventricular zone numbers throughout development relative to control assembloids. Control, n=7 and DS, n=7 for day 30; Control, n=10 and DS, n=9 for day 60; Control, n=9 and DS, n=12 for day 90. Each point on the graph represents one assembloid. (E–G) Quantification of ventricular zone thickness at day 30 (E), day 60 (F), and day 90 (G). No significant difference in ventricular zone thickness was observed at day 30; however, DS assembloids exhibited significantly thinner ventricular zones at day 60 and day 90 compared with controls. Control, n=65 and DS, n=37 for day 30; Control, n=34 and DS, n=8 for day 60; Control, n=17 and DS, n=4 for day 90. Each point on the graph represents one ventricular zone. Data are Mean ± S.D., ns: no significant changes observed, *P < 0.05, **P < 0.01- Welch's t test.

**Figure S10. Neural progenitor depletion in DS assembloids is not associated with increased apoptosis.** (A, C) Representative immunofluorescence images of day 30 and day 90 control and DS assembloids co-stained for SOX2 (green) and the apoptotic marker cleaved caspase-3 (CC3; magenta). (B, D) Quantification of $CC3^+/SOX2^+$ neural progenitors in the hCO and hMGEO regions at day 30 and day 90. No significant differences in progenitor apoptosis were detected between control and DS assembloids in either forebrain compartment. Control and DS, n=5 each genotype for hCO and hMGEO at day 30. Control, n=6 for hCO and hMGEO at day 60; DS, n=4 for hCO and hMGEO at day 90. Each point on the graphs represents one assembloid. Data are Mean ± S.D., ns: no significant changes observed- Welch's t test.

**Figure S11. Increased neuronal population in an independent Dravet syndrome assembloid line.** (A) Representative immunofluorescence images of day 60 assembloids generated from control_2 and DS_2, stained for NeuN (green) and Hoechst (magenta). DS_2 assembloids exhibit an increased abundance of $NeuN^+$ neurons compared with control_2 assembloids. (B) Quantification of $NeuN^+$ cells in the hCO and hMGEO regions at day 60. DS_2 assembloids display a significant increase in the percentage of $NeuN^+$ cells (out of total $Hoechst^+$ cells) in both forebrain compartments relative to control_2 assembloids, independently validating the enhanced neurogenesis phenotype observed in the primary DS patient line. Control_2, n=6 for hCO and hMGEO; DS_2, n=3 for hCO and hMGEO. Each point on the graph represents one assembloid. Data are Mean ± S.D., $^*P < 0.05$, $^{**}P < 0.01$- Welch's t test.

**Figure S12. Inhibitory neuron populations are unaffected in DS forebrain assembloids.** (A) Representative immunofluorescence images of day 90 control and DS forebrain assembloids stained for GABA (green) and Hoechst (magenta). Comparable distributions of $GABA^+$ are observed in both control and DS assembloids. (B) Quantification of $GABA^+$ cells in the hCO and hMGEO regions at day 90. No significant differences in the percentage of $GABA^+$ cells (out of total $Hoechst^+$ cells) were detected between control and DS assembloids in either forebrain compartment. Control, n=6 for hCO and hMGEO; DS, n=4 for hCO and hMGEO. Each point on the graph represents one assembloid. Data are Mean ± S.D., ns: no significant changes observed- Welch's t test.

**Figure S13. Interburst LFP power.** Estimated group effects for interburst absolute (A) and relative (B) LFP power across frequency bands at day 90. Interburst periods were defined as intervals outside detected network bursts in multi. (A) DS assembloids showed higher absolute delta-band power, while other frequency bands did not differ significantly between groups. (B) DS assembloids showed higher relative delta power and lower relative alpha, beta, low-gamma, and high-gamma power. Relative theta power did not differ significantly. Frequency bands were defined as delta, 1–4 Hz; theta, 4–8 Hz; alpha, 8–12 Hz; beta, 12–30 Hz; low-gamma, 30–55 Hz; and high-gamma, 65–100 Hz. Points represent estimated DS minus control effects (beta coefficients). Error bars represent 95% confidence intervals, and the dotted line indicates no group difference. Group differences were assessed using linear mixed-effects models with group as a fixed effect and assembloid identity as a random intercept. When model fitting failed or yielded singular fits, Welch's t-tests were performed on assembloid-level means. Control, n=10; DS, n=10. Asterisks indicate FDR-corrected significant group comparisons (Q < 0.05) after Benjamini-Hochberg correction within the relevant metric family; ns, no significant changes.

**Figure S14. Spatial organization and distance dependence of high-gamma coherence.** Pairwise magnitude-squared coherence was calculated between recording channels in the high-gamma band (65-100 Hz) using artifact-free 1-s Hann windows with 50% overlap. (A) Spatial representation of the 64-channel probe geometry. Each node represents a recording contact, and connecting lines represent pairwise coherence strength, with darker, thicker lines indicating higher coherence. Data showing one representative recording. (B) Difference map of group-average pairwise coherence, calculated as DS minus control. Blue values indicate lower coherence and red values indicate higher coherence in DS assembloids. Channel numbering follows the probe geometry shown in A. (C) High-gamma coherence as a function of interelectrode distance, summarized in fixed 100-μm bins. Dots and lines represent group means across assembloid-level bin means; shaded regions indicate 95% confidence intervals. Coherence varied modestly with interelectrode distance in both groups, with no significant overall group effect or group-by-distance interaction. Coherence-distance relationships were analyzed using a linear mixed-effects model of logit-transformed coherence, with group, interelectrode distance, and their interaction as fixed effects and assembloid identity as a random intercept. Control, n=10; DS, n=10.

**Figure S15. Spike sorting metrics.** (A) Population spike raster from one control (15 units). (B-D) Estimated group effects for single-unit spike-sorting measures in control and DS assembloids. (B) Unit firing and burstiness measures, including spike count per unit, mean firing rate, interspike-interval coefficient of variation, i.e., ISI CV, relative difference in inter-spike intervals between adjacent spikes, i.e., ISI CV2, and within-unit burst fraction. DS assembloids showed a lower within-unit burst fraction than controls; all other measures were not significantly different. (C) Unit yield and population measures, including curated units per recording, active-channel fraction, and sorted-population firing rate. None of these measures differed significantly between groups. (D) Waveform and template-spatial measures, including median extracellular amplitude, template trough-to-peak duration, and template spatial spread. None of these measures differed significantly between groups. Points represent estimated DS minus control effects (beta coefficients), error bars represent 95% confidence intervals, and the dotted line indicates no group difference. Control, n=10; DS, n=10 for panels B-D. Group differences were assessed using linear mixed-effects models with group as a fixed effect and assembloid identity as a random intercept. When model fitting failed or yielded singular fits, Welch's t-tests were performed on assembloid-level means. (E) Signed pairwise CCH map from the same recording as A. Red edges denote pairwise CCH maxima at or above the pooled $75^{th}$ percentile across control and DS assembloids, blue edges denote pairwise CCH minima at or below the pooled $25^{th}$ percentile, and dashed edges indicate that the selected extremum occurred at zero lag. Arrowheads indicate lag order rather than causal direction; node and edge sizes reflect firing rate and CCH magnitude, respectively. Edge color in E denotes the sign of the selected CCH extremum and does not establish excitatory or inhibitory neuronal identity, direct connectivity, or monosynaptic coupling. (F) Mean positive CCH peak. Data are Mean ± S.E.M., ns: no significant changes-Welch's t test. (G) Mean negative CCH trough. Values in G are displayed on a $\times 10^{-3}$ scale. Data are Mean ± S.E.M., ns: no significant changes-Welch's t test. Control, n = 9; DS, n = 5 for panels F and G.

**Figure S16. scRNAseq analysis at day 90.** Integrated UMAP split by sample displaying cell type composition across control and DS assembloids; control = 2 replicates, control_3 = 2 replicates, DS = 2 replicates, DS_2 = 1 replicate.

**Figure S17. scRNAseq analysis at day 90.** Dot plot showing expression of cell type-specific marker genes. CH, cortical hem; ChP, choroid plexus; IPC, intermediate progenitor cell; OPC, oligodendrocyte precursor cell; RGC, radial glial cell.

**Figure S18. Day 30 forebrain assembloids are enriched for neural progenitors and contain few differentiated neurons.** (A) Representative whole-section immunofluorescence images of day 30 control assembloids stained for SOX2 (green), NeuN (magenta), and Hoechst (blue). $SOX2^+$ cells are more abundant, whereas $NeuN^+$ neurons are sparsely detected. (B) Quantification of $SOX2^+$ and $NeuN^+$ cells in day 30 control assembloids. $SOX2^+$ neural progenitors constitute the majority of the cellular population, while $NeuN^+$ differentiated neurons account for a minor fraction of cells. These data confirm that day 30 assembloids represent a progenitor-enriched developmental stage suitable for investigating progenitor calcium-network activity. n=4 for each cell type. Each point on the graph represents one assembloid. Data are Mean ± S.D., **P < 0.01- Welch's t test.

**Figure S19. Intrinsic electrophysiological properties of progenitor-enriched assembloids are unaffacted following *SCN1A* haploinsufficiency.** (A) Section of a day 30 control assembloid immunostained for SOX2 (red) and co-labeled for Biocytin (green). White arrowhead points to the patched $Biocytin^+$ cell which is also $SOX2^+$ (neural progenitor). (B) $C_m$ measured by whole-cell patch-clamp recordings from cells in day 30 control and DS assembloids. No significant differences in membrane capacitance were observed between groups, indicating comparable cellular membrane properties. (C) $R_m$ of recorded cells in day 30 control and DS assembloids. Membrane resistance was not significantly altered by *SCN1A* haploinsufficiency. (D) Holding current measured during whole-cell recordings from control and DS assembloids. No significant differences in holding current were detected between groups. Control, n=9; DS, n=7. Individual cells are represented by points. Data are Mean ± S.D., ns: no significant changes observed- Welch's t test. $C_m$, membrane capacitance; $R_m$, membrane resistance.

**Figure S20. Representative processing of extracellular field recordings for local field potential and multiunit activity analyses.** Representative extracellular recording obtained from a day 90 assembloid. The raw trace (left) illustrates spontaneous network activity recorded using the multichannel neural probe (one of 64 channels shown). The recording was subsequently bandpass filtered to isolate

LFP within the delta (1-4 Hz), theta (4-8 Hz), alpha (8-13 Hz), beta (13-30 Hz), low-gamma (30-55 Hz), and high-gamma (65-100 Hz) frequency ranges (right). MUA was extracted from the same recording, with detected spike events indicated by red markers. These processed signals were used for downstream analyses of spectral power, phase-locking, coherence, burst activity, and spike-train dynamics in control and DS assembloids. LFP, local field potential; MUA, multiunit activity.

**Figure S21. Robustness of sustained calcium-state classification.** (A) Representative ROI-wise ΔF/F traces from one control assembloid and one DS assembloid shown in separate rows for Gaussian smoothing sigma values of 1, 2 and 3 s. Raw traces, smoothed traces and detected active periods are shown; sigma=2 s is the primary setting. (B) Occupancy of the low, middle and large population states across sigma values. Thin lines connect repeated measurements from the same organoid; symbols show individual organoids and group Mean ± S.E.M.; the rightmost panel shows Hedges' g for the DS−control contrast. Low occupancy remained greater in DS assembloids at all three sigma values (DS−control difference, 0.154–0.190; Hedges' g, 1.08–1.57), whereas middle occupancy remained greater in control assembloids (difference, −0.266 to −0.241; Hedges' g, −1.30 to −1.00); the DS-greater large-occupancy difference was smaller. (C) DS−control occupancy differences across the prespecified Schmitt-trigger ON/OFF percentile combinations. The expected low-occupancy direction was retained in eight of nine combinations and the control-greater middle-occupancy direction in all nine combinations. The only low-occupancy direction reversal occurred at the stringent ON/OFF=85/70 setting and was small (DS−control difference, −0.0123; Hedges' g, −0.206). (D) DS−control occupancy differences across the prespecified gap-fill × minimum-duration combinations. Both expected directions were retained in all nine combinations, with tightly stable low differences (0.169–0.171) and middle differences (−0.251 to −0.249). (E) DS−control occupancy differences across the prespecified low-boundary × large-boundary combinations. Both expected directions were retained in all nine combinations; the low difference was attenuated, but remained positive, at the 0.20 low boundary (overall range, 0.100–0.171), while the middle difference ranged from −0.261 to −0.181. (F) Hedges' g across the 27-condition joint sigma × minimum-duration × low-boundary analysis for low and middle occupancy. Both expected effect directions were retained across all 27 joint combinations (low Hedges' g, 0.827–1.62; middle Hedges' g, −1.36 to −0.876). Heatmap color scales are shared within directly comparable panels; outlined cells mark the primary parameter setting. Across the 57 prespecified settings, the DS−control difference in low occupancy remained positive in 56 of 57 settings (98.2%), whereas middle occupancy was consistently greater in control assembloids in all 57 settings. Thus, the low- and middle-occupancy differences were broadly robust to the tested analysis parameters, with a limited exception at the most stringent Schmitt-trigger setting. All group-level summaries use the assembloid as the independent biological replicate (control, n=5; DS, n=9; one recording per assembloid). Exact Welch's P values are reported in the associated source-data tables and were not used to select parameter settings.

### Supplementary figures

#### Figure S1

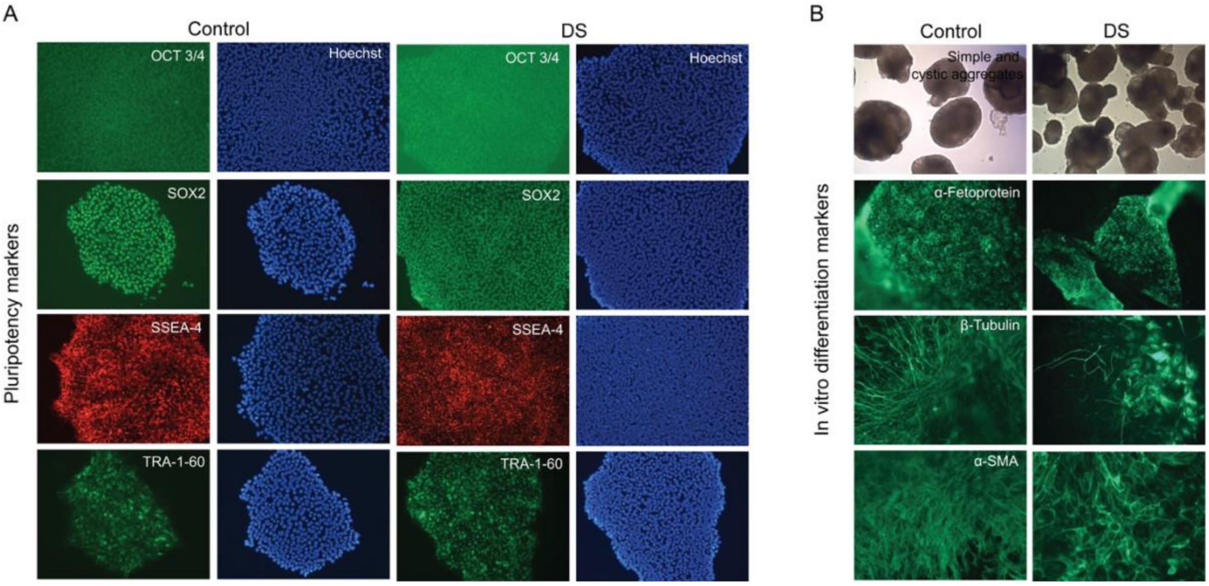


#### Figure S2

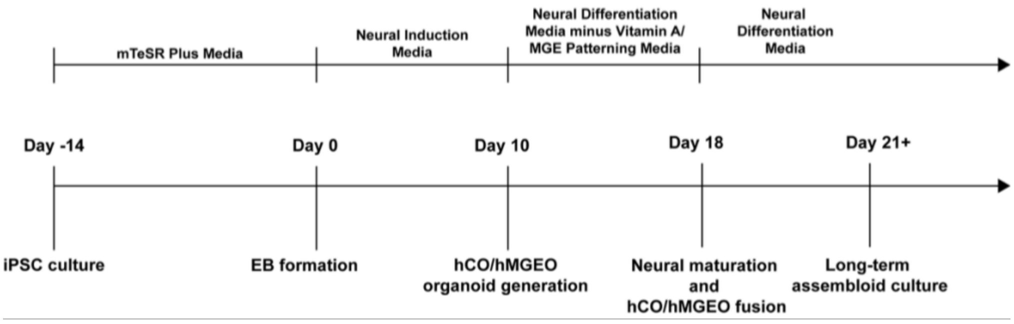


#### Figure S3

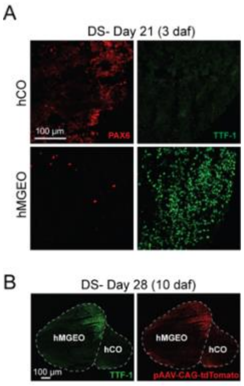

Figure S4

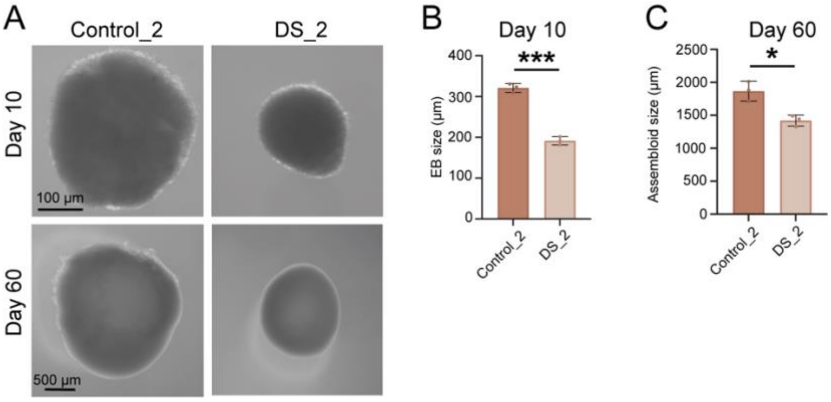


Figure S5

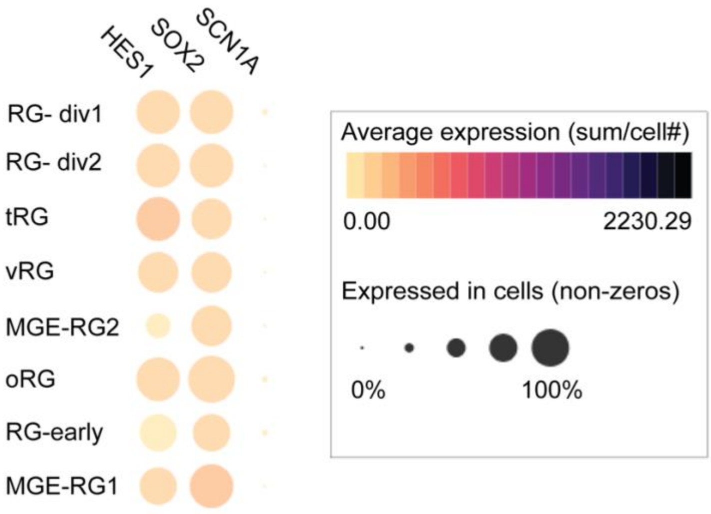


Figure S6

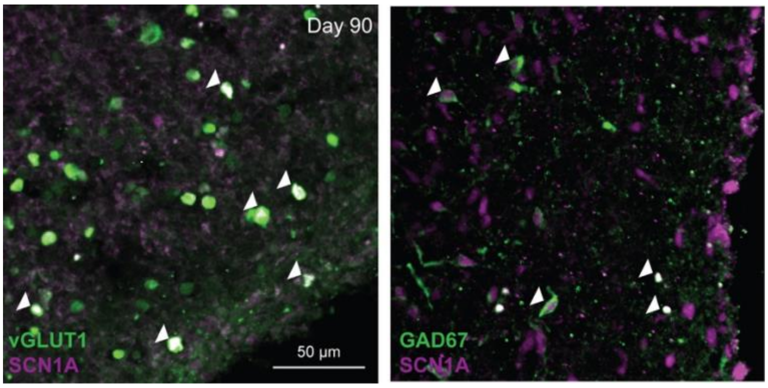

Figure S7

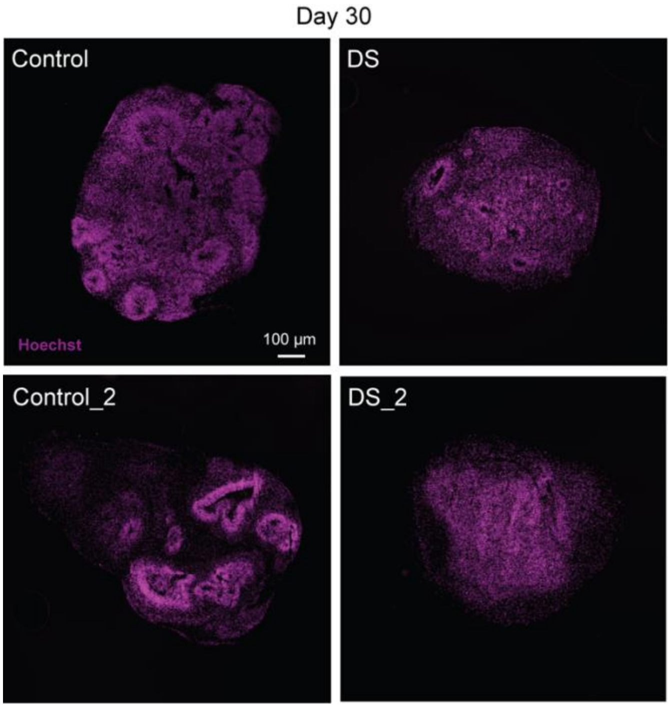


Figure S8

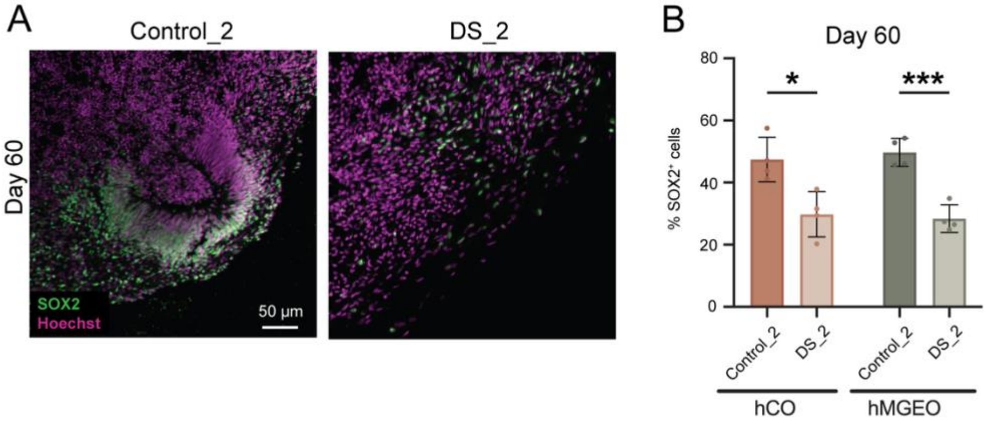

Figure S9

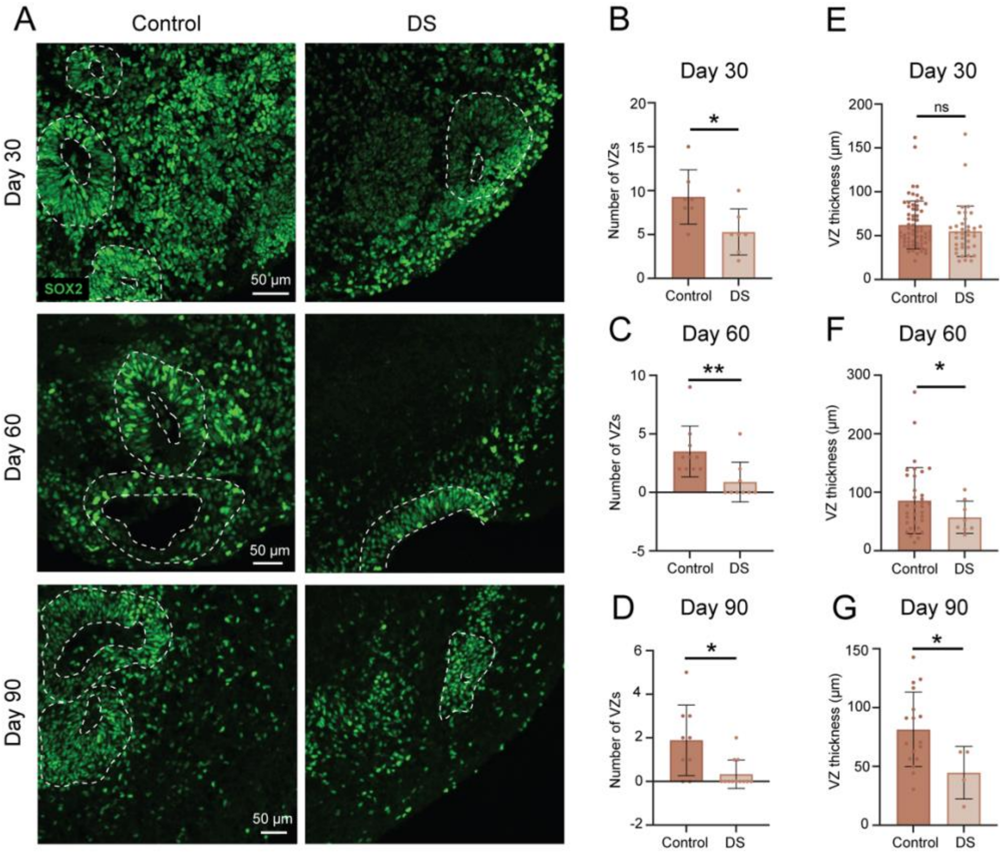


Figure S10

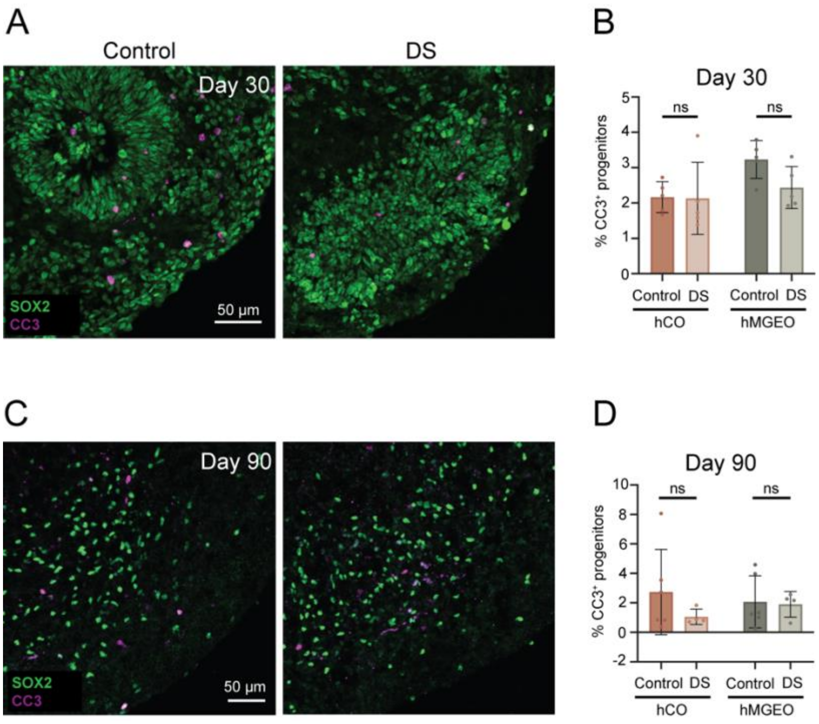

Figure S11

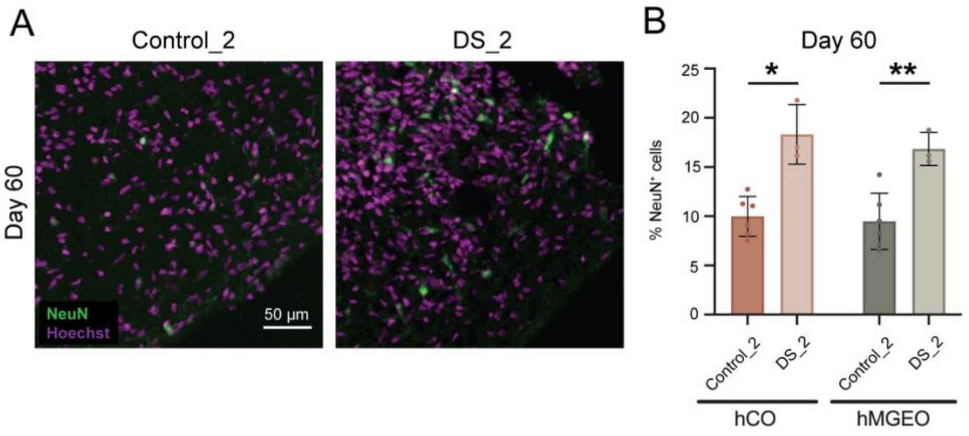


Figure S12

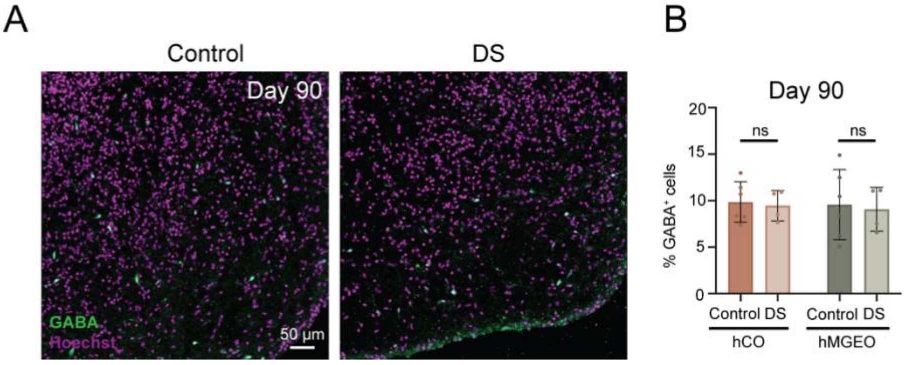


Figure S13

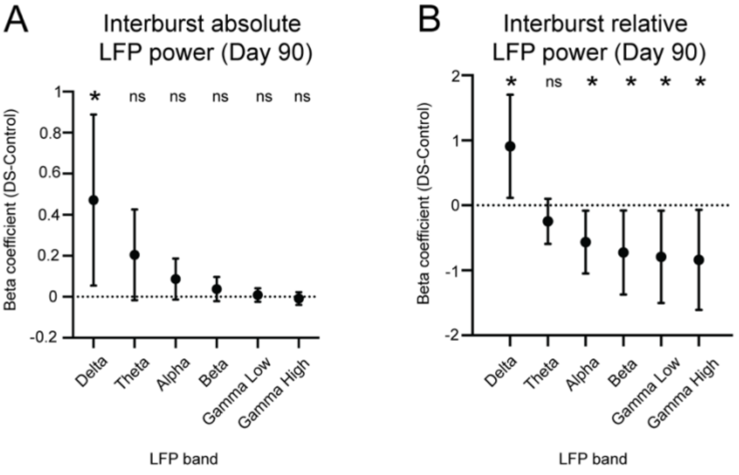

Figure S14

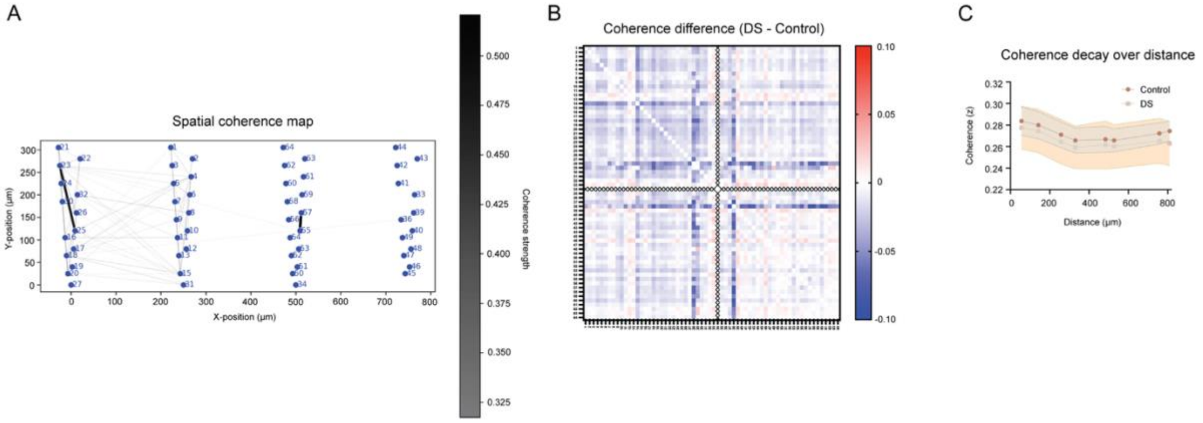


Figure S15

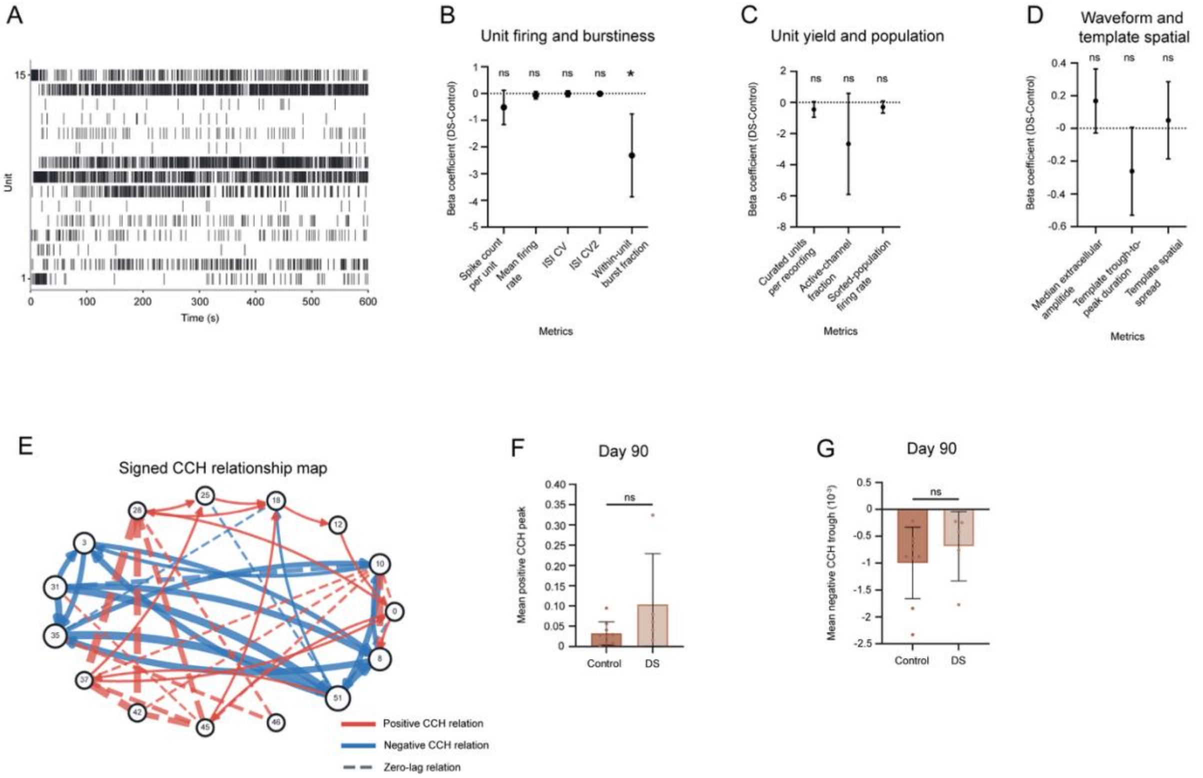


Figure S16

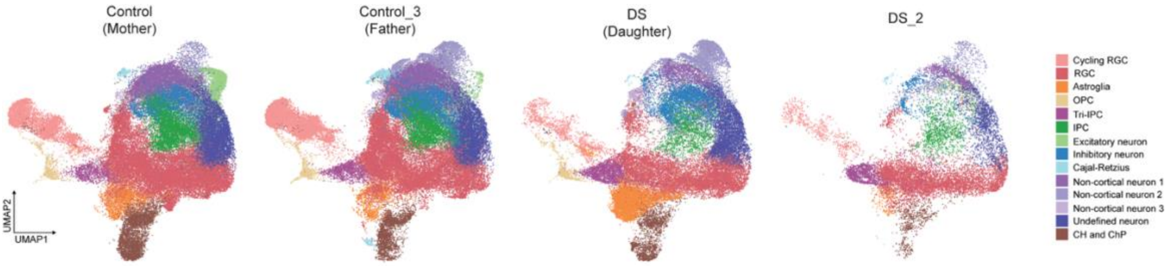

Figure S17

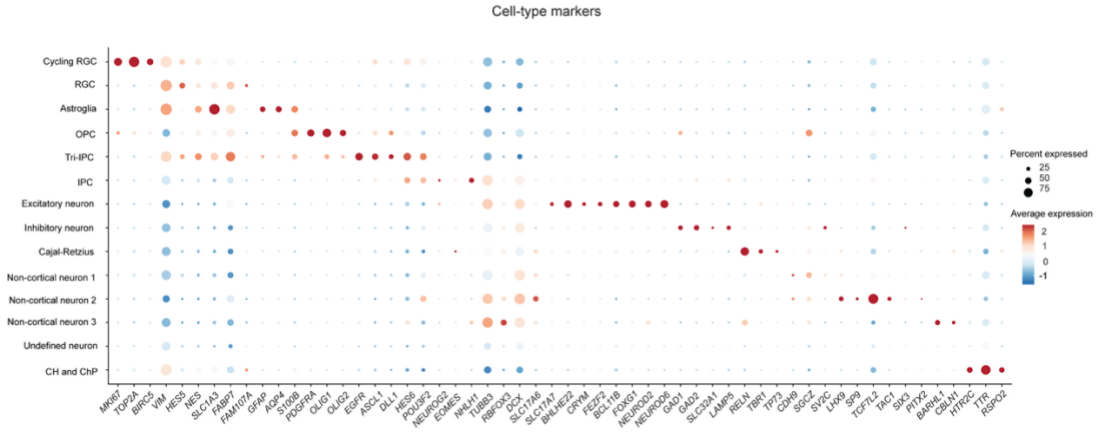


Figure S18

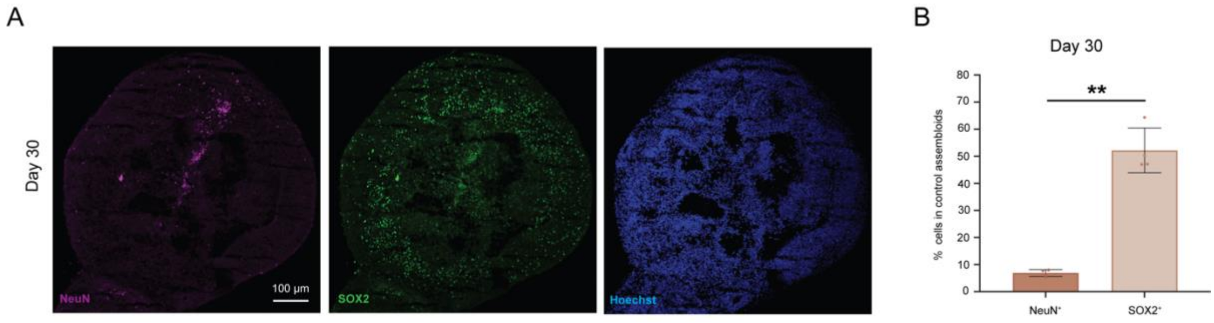


Figure S19

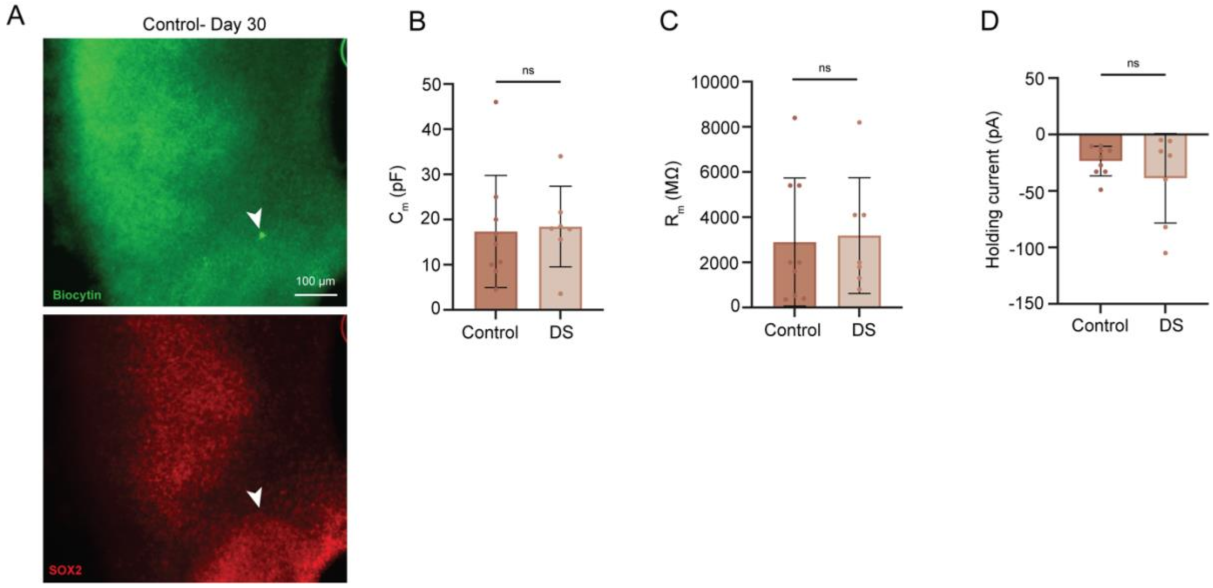

Figure S20

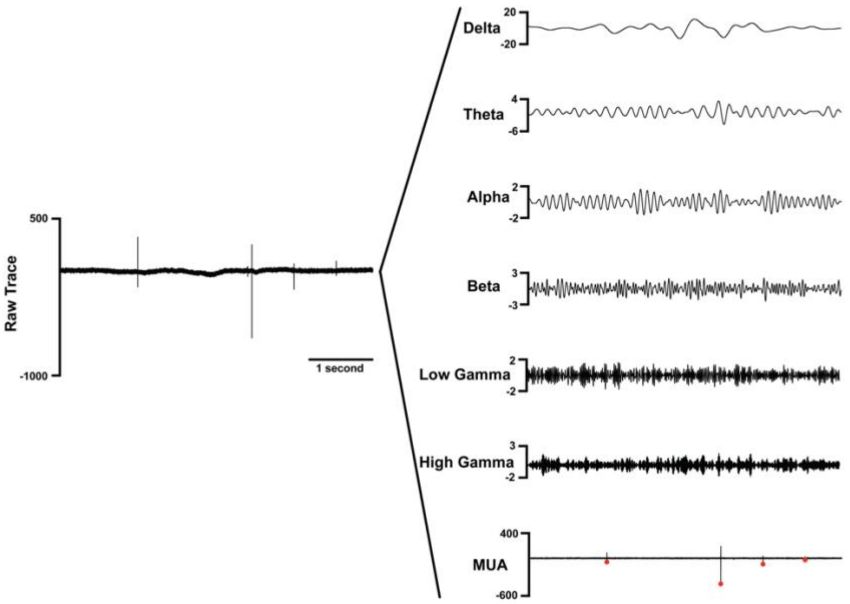


Figure S21

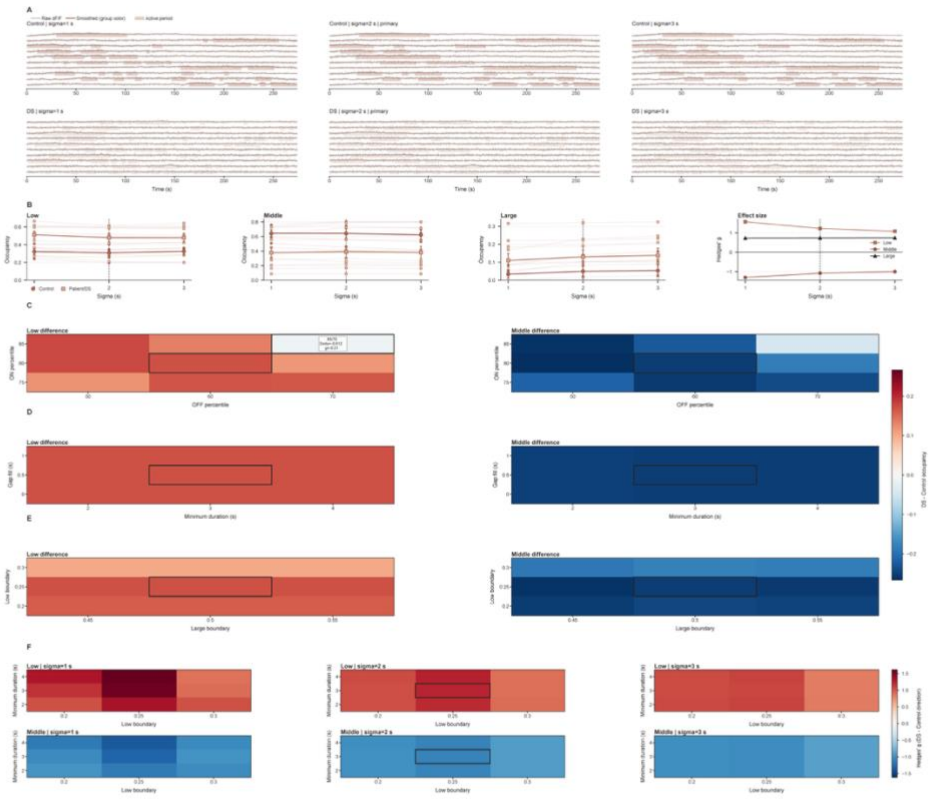

### Supplementary table

**Table S1.** List of antibodies used in immunofluorescence

| Antibody | Supplier | Concentration used |
|---|---|---|
| Anti-TRA-1-160 | Invitrogen (A25617) | 1.2:100 |
| Anti-SSEA4 | Sigma (60062PE) | 1:100 |
| Anti-OCT4 | Sigma (FCMAB113A4) | 1:100 |
| Anti-SOX2 (for pluripotency test) | Sigma (AB5603A4) | 0.5:100 |
| Anti-a-Fetoprotein | Sigma (A8452) | 1:2000 |
| Anti-βIII-tubulin | Sigma (T8578) | 1:2000 |
| Anti- α-SMA | Sigma (A2547) | 1:2000 |
| Anti-PAX6 | Biolegend (901301) | 1:250 |
| Anti-TTF-1 | Santa Cruz Biotechnology (sc-53136) | 1:250 |
| Anti-Calbindin | sWant (CB38) | 1:500 |
| Anti-SCN1A | Abcam (ab24820) | 1:250 |
| Anti-SOX2 | Santa Cruz Biotechnology (sc-365823) | 1:500 |
| Anti-SOX2 (for co-staining with Anti-Ki67 and Anti-NeuN) | EMD Millipore (AB5603) | 1:500 |
| Anti-ZO-1 | Invitrogen (33-9100) | 1:500 |
| Anti-Ki67 | BD Pharmingen (550609) | 1:250 |
| Anti-NeuN | Abcam (ab104224) | 1:500 |
| Anti-vGLUT1 | MilliporeSigma (AB5905) | 1:500 |
| Anti-GAD67 | MilliporeSigma (MAB5406) | 1:500 |
| Anti-CC3 | Cell Signaling Technology (9661) | 1:500 |
| Anti-GABA | MilliporeSigma (A2052) | 1:500 |

### Supplementary movies

**Movie S1.** Live calcium imaging of day 30 control assembloid.

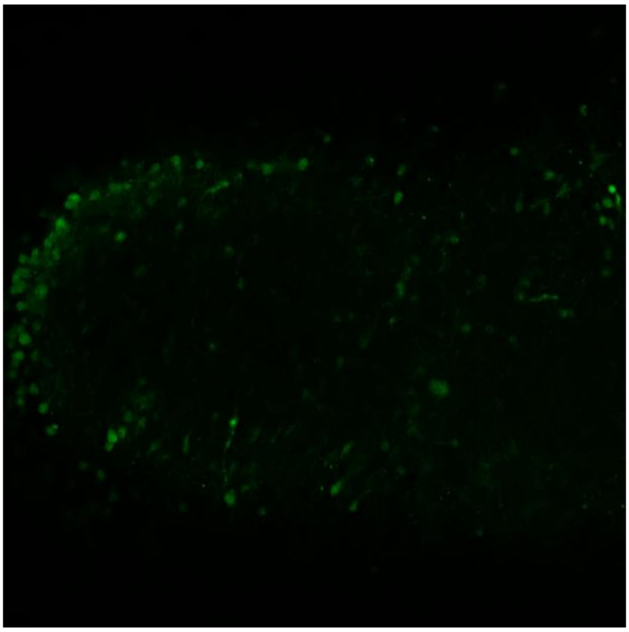

**Movie S2.** Live calcium imaging of day 30 DS assembloid.

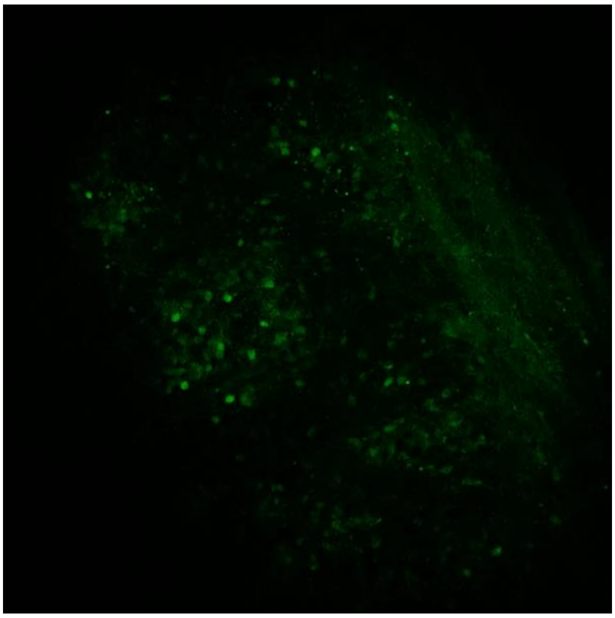
